# Interrogating endothelial barrier regulation by temporally resolved kinase network generation

**DOI:** 10.1101/2022.09.19.508598

**Authors:** Ling Wei, Selasi Dankwa, Kamalakannan Vijayan, Joseph D. Smith, Alexis Kaushansky

## Abstract

Vascular leak is a common disease complication, yet the signaling networks regulating barrier integrity are incompletely understood. We developed a novel methodology, Temporally REsolved KInase Network Generation (TREKING), which combines a 28-kinase inhibitor screen with machine learning and network reconstruction to build time-resolved, functional phosphosignaling networks. We demonstrated the utility of TREKING for identifying pathways mediating barrier integrity following thrombin stimulation with or without tumor necrosis factor (TNF) activation in brain endothelial cells. TREKING assigned distinct barrier phenotypes to mitogen-activated protein kinase (MAPK) pathways and revealed a condition-specific MAPKAPK2/MK2 switch kinase pathway with early barrier-disruptive activity in both inflammatory conditions, but late barrier-restorative activity exclusively in the absence of TNF pre-conditioning. MAPKAPK2/MK2 was activated with expected distinct kinetics under the two inflammatory conditions and late activation was linked to a MAP3K20/ZAK-MAPK14/p38α-MAPKAPK2/MK2 pathway. Beyond MAPKs, TREKING predicts extensive interconnected networks that control barrier integrity and is a tool for dissecting complex temporal phosphosignaling networks across biological systems.

## 1 Introduction

Dysregulation of the blood-brain barrier (BBB) is associated with the pathogenesis of a variety of diseases, including infectious causes such as cerebral malaria and non-infectious such as multifactorial neurodegenerative diseases (Miller et al., 2013, Zhao et al., 2015, Daneman and Prat, 2015). The permeability of endothelial barriers is tightly regulated via cell-cell junctions and focal adhesions. These points of cell-cell and cell-substrate contact are controlled by phosphorylation and actomyosin-mediated contractile mechanisms, whose functions are heavily regulated by diverse phosphosignaling pathways (Mehta and Malik, 2006). Kinases and phosphatases can rapidly (on the scale of seconds to minutes) alter phosphorylation states in response to barrier-disruptive stimuli and can promote long-term regulatory functions during barrier recovery (Kuppers et al., 2014). Whereas kinase signaling pathways mediating barrier function have been intensively explored, there is an incomplete understanding of the temporal regulation of these pathways and their complex network-level connectivity (Kuppers et al., 2014, Komarova et al., 2017, Dankwa et al., 2021, Mehta and Malik, 2006). Moreover, it is difficult to assign specific barrier activity to individual kinases because some kinases are activated in response to both barrier-strengthening and barrier-weakening mediators and/or display distinct time-dependent barrier functions after activation (Birukova et al., 2013, Han et al., 2013, Klomp et al., 2019, Garcia et al., 2001, McVerry and Garcia, 2004, Vouret-Craviari et al., 2002, Knezevic et al., 2009). Thus, understanding the temporal dynamics of barrier regulation and crosstalk between pathways is critical for designing successful targeted therapeutic interventions.

During infection or disease, endothelial cells must integrate complex inflammatory signals. For instance, thrombin and the proinflammatory cytokine tumor necrosis factor (TNF) can both promote morphological remodeling of the endothelium that leads to the loss of barrier integrity (Oldenburg and de Rooij, 2014, Marcos-Ramiro et al., 2014). TNF exacerbates thrombin-induced barrier disruption (Liu et al., 2004, Anrather et al., 1997, Tiruppathi et al., 2001), which indicates that endothelial signaling pathways and/or their kinetics are altered by combined inflammatory stimuli. High-resolution and time-resolved phosphoproteomic analyses have cataloged phosphorylation events that occur in response to thrombin or TNF-induced endothelial activation (van den Biggelaar et al., 2014, Beguin et al., 2019). Separate efforts have developed computational tools and modeling approaches that use-omics data for network generation, including logic modeling such as Boolean logic models and logic ordinary differential equations (ODEs), and enrichment-based approaches such as kinase-substrate enrichment analysis (KSEA) (Casado et al., 2013, Schafer et al., 2019, Vaga et al., 2014). However, these computational tools rely on large-scale phosphoproteomics data that can typically not be collected with fine temporal resolution. Furthermore, proteomics only indirectly assesses kinase functionality, as inferred by phosphorylation states, and many kinases play multiple roles upon activation. Given these limitations, existing network-generation tools have not yet been applied to elucidating the time-resolved molecular functional underpinnings of endothelial barrier integrity.

Kinase regression (KiR) is an approach based on a compound screen that utilizes a small panel of kinase inhibitors with overlapping target specificities and implements elastic net regression to broadly interrogate the function of hundreds of kinases in a cellular phenotype (Gujral et al., 2014, Arang et al., 2017, Dankwa et al., 2021). Kinase inhibitors can alter the temporal features of endothelial barrier disruption and barrier recovery phases following thrombin stimulation (Dankwa et al., 2021), suggesting that different sets of kinases are functional at different stages of barrier perturbation. Here, we develop a novel methodology, Temporally REsolved KInase Network Generation (TREKING), and demonstrate its utility to investigate time-resolved kinase functionality associated with barrier regulation of human brain microvascular endothelial cells (HBMECs) and to build phosphosignaling network models of endothelial barrier regulation.

## 2 Materials and methods

### 2.1 Cell lines

Primary human brain microvascular endothelial cells (HBMECs; Cell Systems Cat# ACBRI 376) were cultured on rat tail collagen type I (5 µg/cm^2^; Corning Cat# 354236) in HBMEC culture media (Lonza Cat# CC-3202) at 37°C and 5% CO_2_. HBMECs were obtained at passage 3 and used until passage 9.

### 2.2 xCELLigence data acquisition

The kinetic data in this study are published and were acquired from xCELLigence assays using the xCELLigence Real Time Cell Analysis (RTCA) Single Plate device (Agilent Technologies) (Dankwa et al. 2021). The assays are described here in brief. HBMECs were grown to confluency in xCELLigence 96 PET E-plates (Agilent Technologies Cat# 300600900). On the day of the assay, HBMECs were equilibrated in serum-free culture media for 1-2 hours, and thrombin (Sigma-Aldrich Cat# T6884) was added at 5 nM. To capture thrombin-induced barrier disruption, the cell index was measured every minute for 6 minutes, after which kinase inhibitors were added in triplicate at 0.5 µM. The cell index was then measured every minute for 2 hours and thereafter, every 5 minutes for 4 hours. The assays with TNF pre-conditioning were performed identically except that HBMECs were activated with 10 ng/ml TNF (R&D Systems Cat# 10291-TA) for ∼22 hours before media equilibration. Data analysis was performed as described previously (Dankwa et al. 2021) with some modifications. The cell index was normalized to baseline (0) at the time point prior to addition of thrombin and area under the curve (AUC) was determined for each kinase inhibitor treatment. For this study, the AUC was determined within 5-minute sliding windows over the 6-hour time course. AUC values were normalized by subtracting the AUC of cells treated with thrombin+DMSO or TNF pre-conditioning +thrombin+DMSO. These values were then linearly transformed, with the most negative normalized AUC value within a 5-minute window being set to 0 and the normalized value for the control sample being set to 100.

### 2.3 Lysate preparation

HBMECs were seeded in 6-well plates (Corning Cat# 353046) at 55,000 cells/well and grown for 3 days. Cells were then activated with 10 ng/ml TNF for 21 hours or kept in media for the same period. On the day of lysate collection, cells were equilibrated in serum-free culture media for 1 hour and then treated with thrombin in triplicate at a final concentration of 5 nM for 5, 15, 30, 60, 120, 180, 240, 360 minutes. After the indicated incubation periods, cells were washed twice with ice-cold phosphate buffered saline (PBS) and lysed in sodium dodecyl sulfate (SDS) lysis buffer (50 mM Tris-HCl, 2% SDS, 5% glycerol, 5 mM ethylenediaminetetraacetic acid, 1 mM sodium fluoride, 10 mM β-glycerophosphate, 1 mM phenylmethylsulfonyl fluoride, 1 mM sodium orthovanadate, 1 mM dithiothreitol, supplemented with a cocktail of protease inhibitors (Roche Cat# 4693159001) and phosphatase inhibitors (Sigma-Aldrich Cat# P5726)). Cell lysates were clarified in filter plates (Pall Cat# 8075) at 3500 rpm for 30 minutes, after which they were stored at −80°C until use. Cell lysates from three biological replicates were collected.

### 2.4 Western blot

All the gel electrophoresis was performed using Bolt™ 4-12% Bis-Tris mini protein gels. Proteins were transferred to PVDF membranes using iBlot 2 dry blotting system (Thermo Fisher Scientific Cat# IB21001). Primary antibodies were used at concentrations recommended by the vendor (see Supplementary Table 3 for antibody information). Antibody to GAPDH (Cell Signaling Technology Cat# 97166, RRID AB_2756824) was used as loading control at 1:2000 dilution. Blots were imaged using Bio-Rad ChemiDoc imaging system and signals were quantified using ImageJ2 (https://imagej.nih.gov/ij/, version 2.3.0). Background correction was done for each band by subtracting background signals nearby the band. The signals from proteins of interest were first normalized to the signals from GAPDH, and then the signals at each time point were normalized to the signals at time zero for the fold change of phosphorylation from basal level. Western blot was performed on cell lysates collected from three biological replicates.

### 2.5 TREKING

To build TREKING models, we incorporated multiple methodologies, the details of which can be found in the methods that follow. Specifically, TREKING models are built by first employing <u>elastic</u> <u>net regularization</u> in a temporally-resolved way that generates a list of predicted functional kinases within each time window. This generates a temporal trace for each predicted kinase, and these temporal traces are then organized into “neurons” using <u>self-organizing map</u> methodology. To generate local networks for the kinases within each “neuron”, we performed <u>network generation</u> steps, which resulted in the comprehensive TREKING models that are described within this manuscript.

#### 2.5.1 Elastic net regularization

The elastic net regularization algorithm used for this study was published previously (Dankwa et al. 2021) but was adapted to generate time windows of kinase activity in barrier function. The kinase regression (KiR) approach exploits the polypharmacology of kinase inhibitors and relies on linear combination of the contributions of kinases to cellular behavior (e.g., barrier permeability in this study) to make predictions on kinases important for specific cell phenotypes and the effect of untested kinase inhibitors on the phenotypes (Gujral et al., 2014). For the temporal KiR (tKiR) approach, a 5-minute sliding time window was applied to the normalized cell index data, which slides at 1-minute steps for the first two hours after kinase inhibitor treatment and at 5-minute steps afterwards. For each time window, the algorithm was applied on the normalized AUC values and kinases with non-zero coefficients were predicted to be informative for barrier function. The sign of the coefficients indicates the functionality of the kinases in regulating barrier integrity, with kinases having positive and negative coefficients being informed to have barrier-strengthening and barrier-weakening function, respectively.

#### 2.5.2 Self-organizing map

To group kinases that were predicted to have similar temporal barrier activity, the NumPy-based SOM implementation MiniSom (https://github.com/JustGlowing/minisom) was used to cluster the tKiR predicted kinases by their temporal characteristics. Grid size of 6×6 was used to generate SOMs. For each SOM neuron, the barrier activity of kinases was plotted over time as a proportion of the kinases in that neuron. For example, if a neuron consisted of 4 kinases and all had the same predicted barrier activity during a specific time window, then they were assigned “+1” (if barrier-strengthening) or “-1” (if barrier-weakening), indicating that 100% of kinases had that same barrier activity. However, if only half of the kinases in that neuron (2/4 kinases) were active during a different time window, then it was assigned +0.5/-0.5, and so on.

#### 2.5.3 Network generation

To build the local phosphosignaling network for an SOM neuron, the shortest paths between any pair of kinases within that neuron was identified using the kinase-substrate phosphorylation database on PhosphoSitePlus^®^ as the background network, and all the paths were combined to form the local phosphosignaling network that describes the paths through which the signals may propagate with certain kinetics. Search for the shortest paths between kinases was done using NetworkX (https://github.com/networkx/networkx), a Python package for analyzing complex network structures.

### 2.6 Literature search

To compare the scope of previous research with the current study, a comprehensive literature search on protein kinases reported to regulate barrier function was done using the free search engine PubMed (https://pubmed.ncbi.nlm.nih.gov). The search terms were “endothelial barrier thrombin” plus kinase gene names, and the search results were filtered so that only articles published on or after year 2000 were shown. The search was done on each of the 518 human protein kinases. The search was not restricted to studies on HBMECs; instead, studies on thrombin-induced barrier disruption using endothelial cell lines, primary endothelial cells isolated from different organs, or in vivo studies using mice or rats, were all included. To compare with our work, studies or data beyond a 6-hour time window (after thrombin treatment) were excluded.

### 2.7 Visualization

Schematics were created with BioRender.com. Phosphosignaling networks were visualized using Cytoscape (https://cytoscape.org, version 3.7.2), with the hierarchic layout from yFiles layout algorithms (https://www.yworks.com/products/yfiles-layout-algorithms-for-cytoscape).

### 2.8 Statistical analysis

Student’s t-test was used to evaluate the difference of kinase activity between non-treated and thrombin-treated conditions. SciPy statistical package (https://github.com/scipy/scipy, version 1.8.0) was used to perform the t-tests. At each time point, both p-value and fold change of phosphorylation with respect to non-treated cells were reported to determine if the two sets of data are different from each other. A p-value below 0.05 or all biological replicates reporting a fold change increasing/decreasing by at least 20% compared to non-treated cells indicates that the kinase activity is different between non-treated and thrombin-treated conditions.

## 3 Results

### 3.1 TNF pre-conditioning alters kinase determinants of thrombin-induced barrier perturbations

To investigate kinase signaling pathways induced by thrombin and explore how the phosphosignaling environment may be altered by sequential inflammatory stimuli, we exploited our published KiR dataset of HBMEC monolayers stimulated with thrombin, with or without TNF pre-conditioning (Dankwa et al., 2021). Thrombin causes an acute drop in HBMEC barrier integrity that peaks around 20-30 minutes and a slower barrier recovery over the next 1.5 hours (Figure 1A, Supplementary Figure 1). TNF pre-conditioning exacerbated thrombin-induced barrier disruption, as demonstrated by the increased maximum barrier permeability and prolonged barrier recovery (Figure 1A, Supplementary Figure 1). While this finding is consistent with previous reports (Tiruppathi et al., 2001, Liu et al., 2004, Anrather et al., 1997), the molecular networks that regulate these differences remain unknown.

**FIGURE 1.**
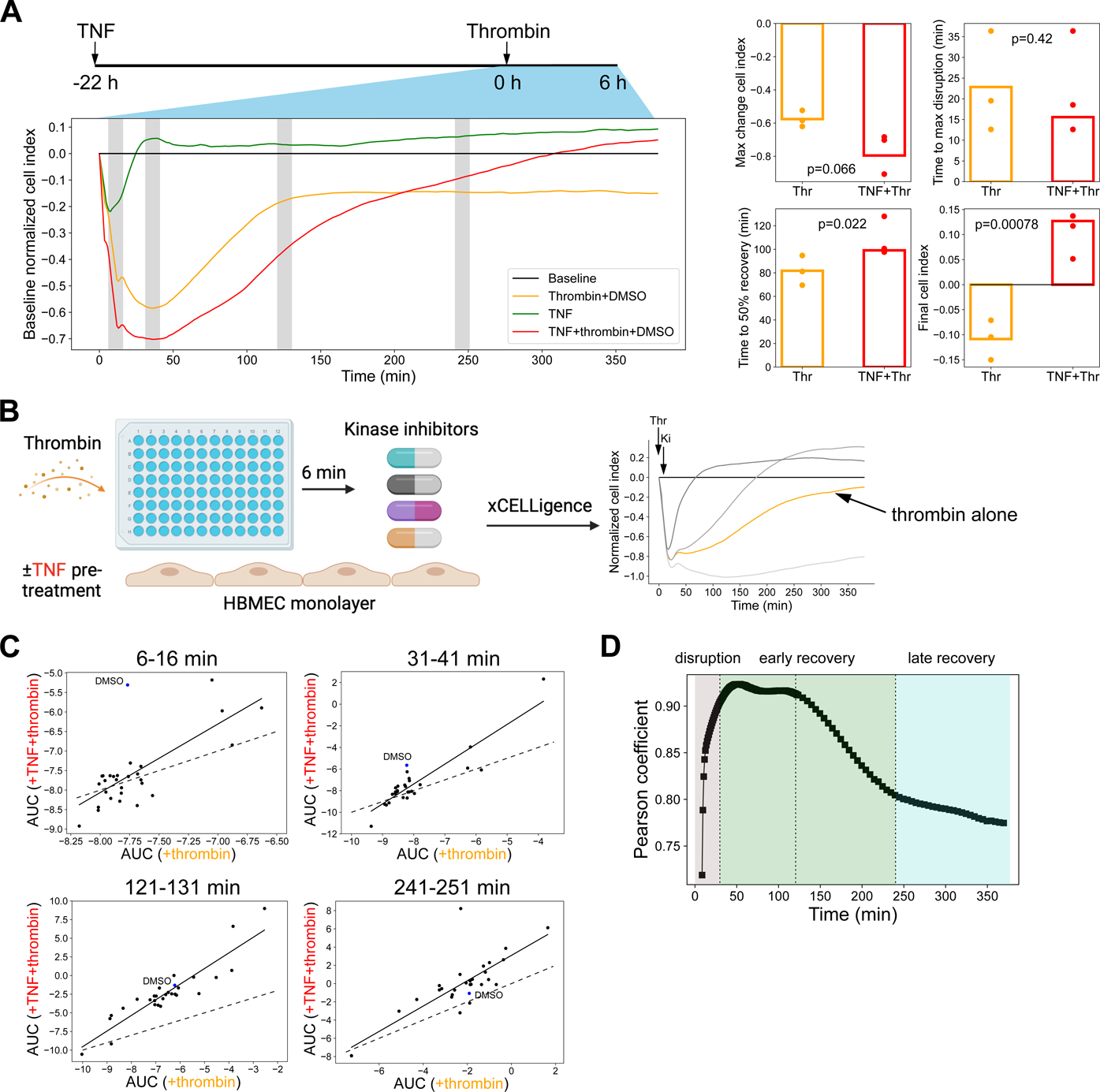
Pre-conditioning with TNF alters the temporal kinetics and magnitude of the thrombin-induced barrier disruption response. **(A)** Left: representative xCELLigence data showing barrier response to thrombin with or without TNF pre-conditioning. Right: Quantification of the maximum change and time to 50% recovery (N = 3). P-values from student’s t-test are shown. **(B)** Graphic of KiR screen conducted with 28 kinase inhibitors for their impact on barrier permeability in thrombin-treated HBMECs (-/+TNF pre-conditioning). Monolayers of HBMECs were treated with thrombin followed at 6 minutes by kinase inhibitor treatment, and barrier permeability was measured in real-time using the xCELLigence system. The xCELLigence curves on right illustrate kinase inhibitor changes in the thrombin-induced barrier response (reprinted from Figure 2F of Dankwa et al., 2021.) **(C)** Correlations of the 28 kinase inhibitors between +thrombin and TNF pre-conditioning +thrombin conditions at different stages of barrier perturbation. The area under the corresponding permeability curve (AUC) within representative time windows (10-minute) is displayed as scatter plots (blue circle: DMSO-treated; black circles: kinase inhibitor-treated; black solid line: linear fitting line excluding DMSO control; black dashed line: identity line). The corresponding time windows to the scatter plots are indicated in panel **(A)** with gray shading. **(D)** Pearson correlation coefficients between AUC in +thrombin and TNF pre-conditioning +thrombin conditions. Temporal AUC was calculated within a 5-minute sliding time window that slides at 1-minute steps for the first 2 hours of kinase inhibitor treatment and at 5-minute steps afterwards. See also Supplementary Figure 1.

To define kinase signaling mechanisms underlying TNF potentiation of thrombin-induced barrier disruption, we reanalyzed a dataset from a small chemical compound screen against human kinases. In the original kinase regression analysis, HBMEC monolayers were treated with thrombin (-/+TNF pre-conditioning) and 28 kinase inhibitors were added six minutes after thrombin treatment (Figure 1B) (Dankwa et al., 2021). The endothelial barrier integrity was assessed in real-time using the xCELLigence system, which measures cellular impedance across adherent cells in electronic microplate wells. Decreases in cell index correlate with the initial strong retraction of endothelial cells following thrombin treatment during the barrier disruption phase, while increased cell index values are associated with recovery of monolayer integrity (Dankwa et al., 2021). We observed that the DMSO vehicle alone had no effect on HBMECs treated with thrombin or TNF (Supplementary Figure 1C), but that different kinase inhibitors blunt, have minimal effect, or exacerbate thrombin-induced barrier disruption and exhibit temporal features during the 6-hour time window (Figure 1B) (Dankwa et al., 2021).

We reasoned that if similar kinases control the barrier integrity in the two inflammatory conditions, then there should be a stronger correlation between the activities of the 28 kinase inhibitors in thrombin alone and TNF pre-conditioned settings. In contrast, if different kinases control the barrier integrity in the two conditions, we would expect less correlation between the activities of the kinase inhibitors. Thus, we reasoned that the KiR screen can be used to probe the underlying barrier regulatory phosphosignaling networks across conditions. Using area under the curve (AUC) of the normalized cell index as the metric, we compared the two inflammatory conditions across four different time windows: barrier disruption (6-16 minutes after thrombin treatment), early barrier recovery (31-41 minutes after thrombin treatment), mid barrier recovery (121-131 minutes after thrombin treatment) and late barrier recovery (241-251 minutes after thrombin treatment) (Figure 1C). We also systematically evaluated the correlation between the activities of the 28 kinase inhibitors in +thrombin and TNF pre-conditioning +thrombin conditions within a 5-minute time window that slides at 1-minute steps for the first two hours after kinase inhibitor treatment and at 5-minute steps afterwards (Figure 1D). The correlation was the lowest in the early barrier disruption phase (Pearson’s r = 0.72) and increased during early barrier recovery (Pearson’s r = 0.90 to 0.92) (Figure 1D), suggesting that the phosphosignaling networks are more divergent during barrier disruption and more similar during the initial barrier restoration phase. Notably, the correlation declined again at approximately 120 minutes and remained divergent throughout the entire mid-to-late barrier recovery phases (Pearson’s r = 0.91 to 0.77), a period when the two inflammatory conditions diverged in their slopes of recovery and final cell index (compare Figure 1A and Figure 1D). This analysis is consistent with a model that TNF pre-conditioning alters kinase regulatory networks that dictate the extent and kinetics of thrombin-induced barrier disruption and recovery in brain endothelial cells.

### 3.2 Time-resolved predictions of barrier-weakening and barrier-strengthening kinases between the two inflammatory conditions

Previously, we predicted 29 and 25 kinases that regulate the HBMEC barrier in response to thrombin treatment with or without TNF pre-conditioning, respectively (Dankwa et al., 2021). However, these KiR predictions were based on the full AUC (6-hour time window, post-thrombin treatment), and did not consider time windows of barrier activity. To address this lack of temporal resolution, we performed temporal KiR (tKiR) using a sliding window analysis (Figure 2, top, schematic overview). For this approach, a 5-minute sliding time window was applied to the normalized cell index data, which slides at 1-minute steps for the first two hours after kinase inhibitor treatment and at 5-minute steps afterwards (Figure 2, Figure 3A). Among the 291 protein kinases used for tKiR predictions (Anastassiadis et al., 2011) (Figure 2, see methodology), the number of functional kinases increased to 120 and 108 in HBMECs treated with thrombin −/+TNF pre-conditioning, respectively (Figure 3A, Supplementary Table 1). The higher resolution of tKiR is because the sliding window analysis allows for the detection of kinases whose activity is brief or sporadic. In total, 159 kinases (∼30% of the total human kinome) were predicted to regulate barrier function within the 6-hour timeframe, with 69 kinases being common to the two conditions (Figure 3A).

**FIGURE 2.**
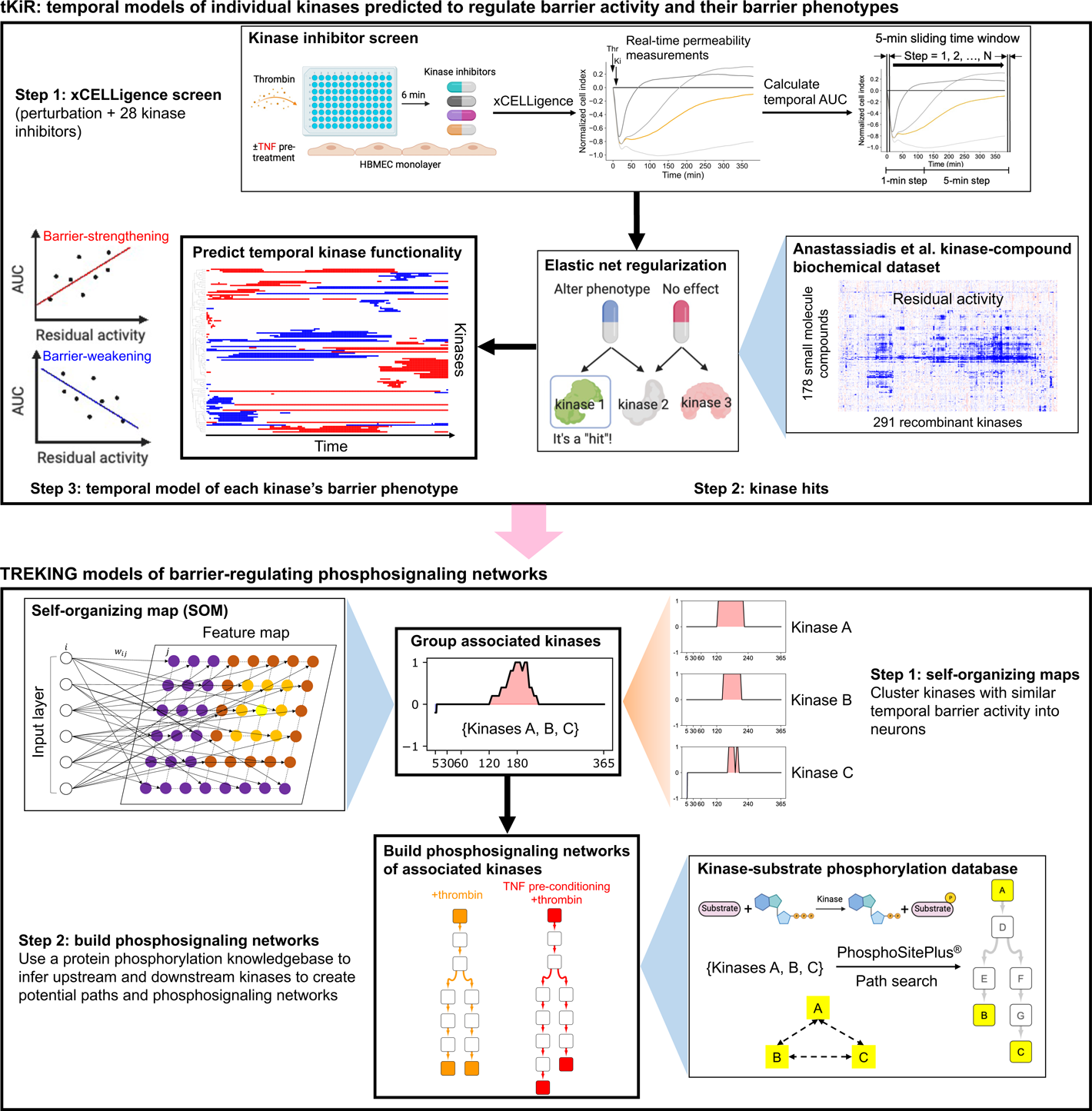
Workflow for building TREKING models from tKiR predictions. Top: tKiR models of temporal kinase barrier functionality was predicted from a 28-kinase inhibitor screen using the elastic net regularization algorithm. In brief, HBMECs were stimulated with thrombin (with or without TNF pre-conditioning) and 28 kinase inhibitors were added 6 minutes after thrombin stimulation to perform kinase regression analysis. The barrier properties of endothelial cells were measured in real-time using the xCELLigence system. The kinase inhibitor screen data (Dankwa et al., 2021) and kinase-compound biochemical data (Anastassiadis et al., 2011) were used for tKiR. Temporal AUC was calculated within a 5-minute sliding time window that slides at 1-minute steps for the first 2 hours of kinase inhibitor treatment and at 5-minute steps afterwards. The barrier function of kinases (barrier-weakening or barrier-strengthening functionality) was predicted at each time window indicated above based on the slope of the 28 kinase inhibitors AUC values versus the residual kinase activity from the Anastassiadis et al dataset. Bottom: workflow for building TREKING networks to describe kinase phosphosignaling in thrombin-activated barriers. Self-organizing maps (SOMs) were used to identify kinases with similar time-resolved barrier activity across the 6-hour timeframe following thrombin treatment. Local phosphosignaling networks for each neuron were built by searching for the shortest paths between any pair of kinases within that neuron using the kinase-substrate phosphorylation database on PhosphoSitePlus^®^ as the background network.

**FIGURE 3.**
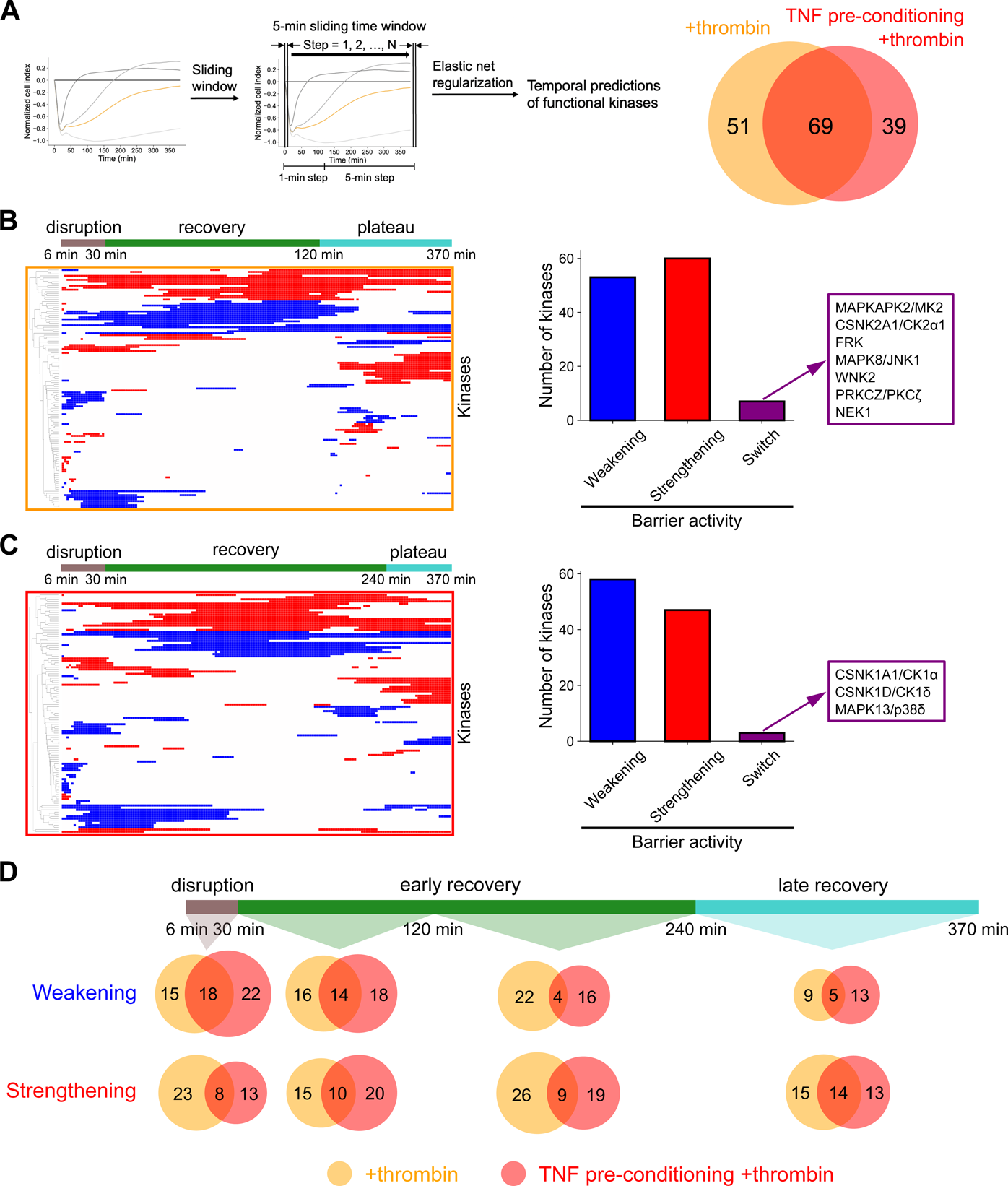
Using kinase regression to inform kinases important for barrier regulation with temporal resolution. **(A)** Left: schematic describing the tKiR methodology to predict time windows of kinase barrier activity. A 5-minute sliding time window was applied to the normalized cell index data from the xCELLigence assays conducted with the 28 kinase inhibitors. Right: Venn diagram showing the total number of kinases predicted in each condition across all time windows. **(B)**-**(C)** Left: time-resolved barrier activity profiles of predicted kinases in **(B)** +thrombin and **(C)** TNF pre-conditioning +thrombin conditions. Blue and red colors represent barrier-weakening and barrier-strengthening functionality, respectively. Predicted kinases are hierarchically clustered using Euclidean distance metric. Right: number of kinases predicted to be barrier-weakening, barrier-strengthening, or of dual functionality across the time course (“switch kinases”). **(D)** Venn diagrams showing the number of kinases predicted to be barrier-weakening or barrier-strengthening at different stages of barrier perturbation. See also Supplementary Table 1.

As a further refinement, tKiR models were used to predict the kinase functionality (promoting barrier weakening or promoting barrier strengthening) in a time-resolved manner (Figure 3B-C). The barrier activity of individual kinases was inferred by the linear regression slope of the kinase inhibitors targeting that kinase from the kinase regression analysis, where barrier-weakening or barrier-strengthening kinases are defined as the kinases whose increased activity leads to barrier disruption or strengthening of barrier integrity, respectively (Figure 2, top). For HBMECs not pre-conditioned with TNF, 53 and 60 kinases were predicted to play a barrier-weakening and barrier-strengthening role, respectively (Figure 3B). In addition, seven kinases were predicted to play both barrier-weakening and barrier-strengthening roles during different time windows and were termed “switch kinases” (Figure 3B). For HBMECs pre-conditioned with TNF, 58 and 47 kinases were predicted respectively to be barrier-weakening and barrier-strengthening, and 3 switch kinases were predicted (Figure 3C). Overall, a greater number of barrier-weakening kinases were predicted during the initial disruption phase, whereas the proportion of barrier-strengthening kinases increased over barrier recovery (Figure 3D). Nevertheless, cells propagate both types of barrier activities concurrently and it is the balance that changes over time (Figure 3D).

### 3.3 The tKiR predictions of kinase activity recapitulate information flow through canonical MAPK signaling pathways involved in barrier regulation

Previous work has established that MAPK cascades are activated after thrombin stimulation, including the extracellular signal-regulated kinases (ERKs), Jun N-terminal kinases (JNKs) and p38 MAPKs (Mehta and Malik, 2006, Radeva and Waschke, 2018, Minami et al., 2004). The topology of MAPKs is well-established; cell surface receptors such as receptor tyrosine kinases (RTKs) provide a link between extracellular stimuli and the cascade (Morrison, 2012). After activation, most MAPK pathways have a four-tier kinase architecture (MAP3K-MAP2K-MAPK-MAPKAPK) (Pimienta and Pascual, 2007). Given the strict directionality of the network, we reasoned that RTKs would be predicted to regulate the barrier during the earliest time points after thrombin treatment, followed sequentially by MAP3Ks, MAP2Ks, MAPKs and then MAPKAPKs. Furthermore, kinases involved in a signaling cascade would be more likely to have the same barrier activity over a given time window, especially if they were acting as relays to propagate a specific barrier phenotype.

Overall, the ERK, JNK, and p38 pathways were all implicated in thrombin signaling, with multiple kinases in each signaling cascade predicted by tKiR (Figure 4). Specifically, within the ERK and the JNK pathways, majority of the predicted kinases were barrier-weakening, and the windows when their activity was predicted followed the expected order of MAP3Ks or MAP2Ks preceding MAPKs, which preceded downstream MAPKAPKs (Figure 4). Whereas the ERK pathway had both an early and a late wave of barrier-weakening activity, the JNK pathway was predicted to have primarily early barrier activity (Figure 4). By comparison, the p38 pathway had a mixture of early barrier-weakening activity and late barrier-strengthening activity, which was mediated by different p38 isoforms (Figure 4). Notably, MAPKAPK2/MK2, a switch kinase, had an early barrier-weakening activity and late barrier-strengthening activity (Figure 4). MAPKAPK2/MK2 is a primary target of MAPK14/p38α, although it can also be activated by MAPK1/ERK2 and JNK signaling (Ronkina et al., 2008, Johnson et al., 2023). Of interest, a mid-to-late acting barrier-strengthening p38 pathway could be assembled from known components, consisting of MAP3K20/ZAK-MAPK14/p38α-MAPKAPK2/MK2, but different upstream kinases appeared to be involved in the early barrier activity of MAPKAPK2/MK2 under thrombin stimulation (Figure 4).

**FIGURE 4.**
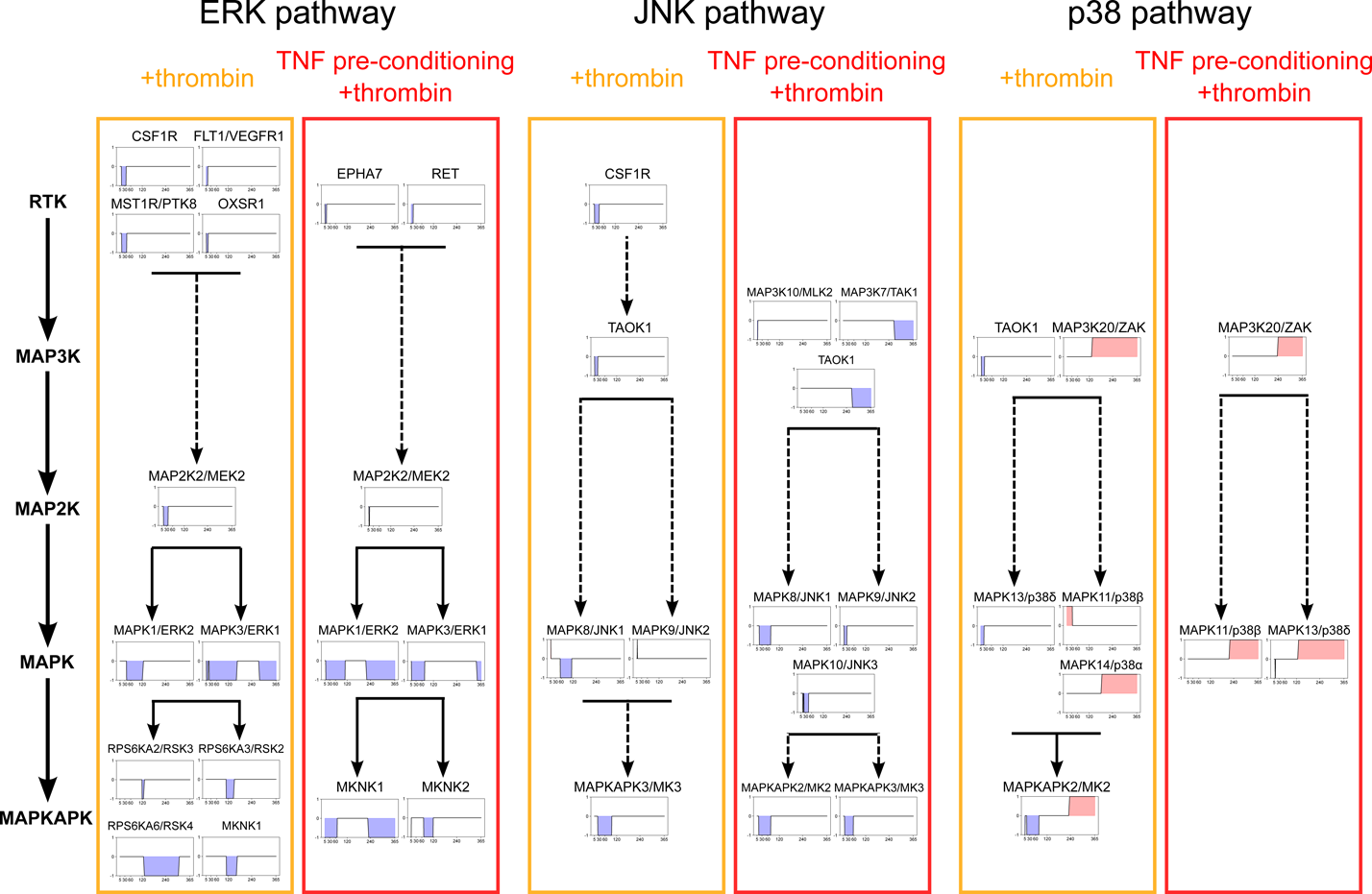
The tKiR methodology recapitulates the phosphosignaling events in canonical MAPK pathways. Members of three canonical MAPK pathways (ERK, JNK, and p38) were predicted to regulate barrier function in thrombin-treated HBMECs with or without TNF pre-conditioning. Kinases were mapped onto the corresponding signaling modules along with their temporal functionality profiles, where blue and red shadings represent barrier-weakening and barrier-strengthening activity, respectively. Solid arrows represent direct kinase phosphorylation interactions reported previously and dashed arrows represent indirect interactions as reported in previous literature (Cargnello and Roux, 2011, Ronkina and Gaestel, 2022). See also Supplementary Table 1.

The same MAPK pathways were predicted with TNF pre-conditioning; however, the timing and duration of the ERK and the JNK pathways were shifted earlier and shortened for most kinases (Figure 4). TNF pre-conditioning also modified barrier-regulatory signaling cascades within the ERK, JNK, and p38 pathways. In the ERK pathway, there was a second wave of barrier-weakening activity by MAPK1/ERK2 and in the JNK pathway, the barrier-weakening activity of TAOK1 was also shifted from an early to a later timeframe. TAOK1 can promote activation of the ERK1/2 pathway in macrophages (Zhu et al., 2020), suggesting that TAOK1 may also function in the ERK pathway in regulating barrier properties. In the JNK pathway, the number of barrier-regulatory kinases was increased, consistent with previous work indicating that combined TNF and thrombin treatment alters the magnitude and duration of JNK activation in endothelial cells (Liu et al., 2004). In the p38 pathway, the barrier activity of the p38 isoforms was rewired. Whereas MAPK13/p38δ retained a brief barrier-weakening activity, the late barrier-strengthening MAPK14/p38α-MAPKAPK2/MK2 pathway was truncated and other p38 isoforms replaced it. Therefore, MAPKAPK2/MK2 was a condition-specific, switch kinase. It had early barrier-weakening activity in both inflammatory conditions, but late barrier-strengthening activity only in the thrombin alone condition. Consequently, this analysis suggests that core MAPK signaling pathways involved in thrombin-induced barrier disruption remain intact after TNF pre-conditioning, but the timing, duration, and importance of specific MAPKs are altered.

### 3.4 Kinase activation state is largely consistent with tKiR predicted barrier activity in the ERK and the JNK pathways

To investigate the tKiR predictions about MAPK signaling pathways regulating barrier phenotypes in the two inflammatory conditions, we probed kinase phosphorylation over time via western blot. We reasoned that there should be a rise in phosphorylation at kinase activation sites immediately preceding and/or during periods of predicted barrier activity. For this analysis, HBMECs (-/+TNF pre-conditioning) were treated with thrombin for 0, 5, 15, 30, 60, 120, 180, 240 or 360 minutes. In each experimental condition, the level of phosphorylated kinase after thrombin treatment was compared to its phosphorylation at the basal level (media only) after normalization to GAPDH. Initially, we probed the ERK and JNK pathways that were primarily predicted to be involved in barrier-disruptive signaling cascades from the tKiR analysis. Control western blots showed that the total levels of the ERK1/2 and JNK1/2 isoforms did not change in this period (Supplementary Figure 2A). Within the canonical ERK cascade, there was sequential activation of kinases, and activation marks appeared in MAP2K1/2 (MEK1/2), MAPK1/ERK2, MAPK3/ERK1, and the downstream MAPKAPK target MKNK1 prior or during predicted early barrier-weakening activity in both inflammatory conditions (Figure 5A). This finding is consistent with the tKiR model (Figure 4). In addition, there was a second wave of MAPK1/ERK2 and MAPK3/ERK1 late barrier activity in both inflammatory conditions that was independent of late MAP2K1/2 (MEK1/2) activation (Figure 5A). As MAP2K1/2 (MEK1/2) were only predicted to have early barrier activity (Figure 4), this raises the possibility that another kinase(s), such as TAOK1, is responsible for the late MAPK1/3 (ERK2/1) activation or there is a change in the activity of a phosphatase(s) that can dephosphorylate MAPK1/3 (ERK2/1).

**FIGURE 5.**
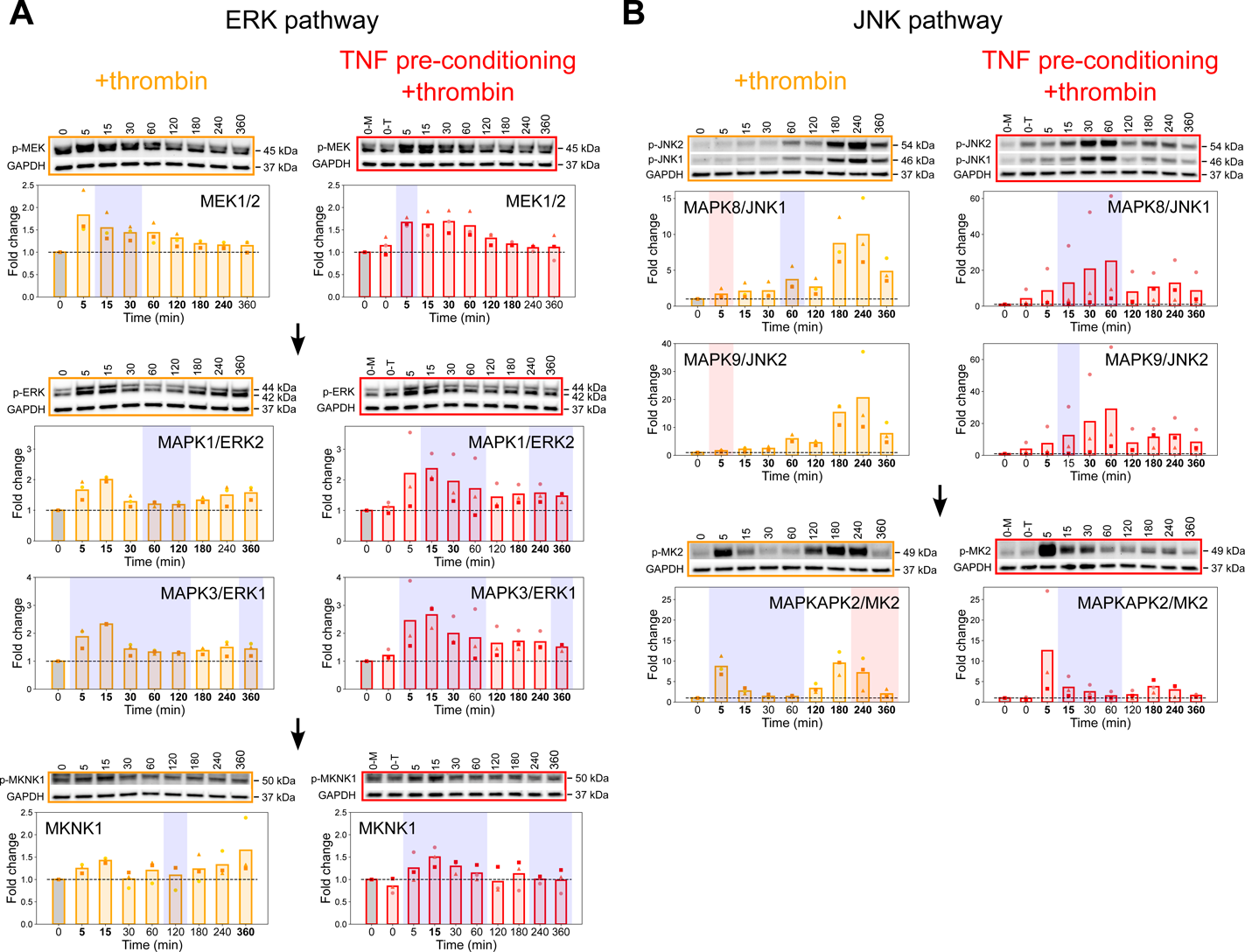
Correlation between phosphorylation activation marks in ERK and JNK kinases and tKiR predicted barrier activity. To study the correlation between kinase activation marks and predicted barrier activity, HBMECs were stimulated with TNF (10 ng/ml) for 21 hours or left unstimulated, and cells were then treated with thrombin (5 nM) for the indicated times. Phosphorylation of members in **(A)** ERK and **(B)** JNK pathways was probed by western blot in three biological replicates. Cells pre-conditioned with TNF but not treated with thrombin are labeled “0-T”. Data was normalized to non-treated, media only condition (“0-M”, gray bars). GAPDH was used as loading control. Symbols represent the fold change of individual biological replicates, and bars represent the mean fold change of three biological replicates. Blue and red shadings represent respectively barrier-weakening and barrier-strengthening activity, as predicted by tKiR. The bolded timepoints indicate that the phosphorylation is different from the basal level (p-value from student’s t-test below 0.05 or all biological replicates reporting a fold change increasing by at least 20% compared to non-treated cells). See also Supplementary Figure 2 and Supplementary Table 3.

Results from quantitative western blot analysis also showed that activation marks increased in both MAPK8/JNK1 and MAPK9/JNK2 prior to predicted early barrier-weakening activity (Figure 5B). Moreover, the timing, magnitude, and duration of early MAPK8/JNK1 activation increased with TNF pre-conditioning, consistent with a shift in the timing of barrier activity predicted by tKiR (Figure 5B). There were also some notable discrepancies between activation marks and predicted barrier activity. In +thrombin condition, we only observed very modest increases in the activity of MAPK8/JNK1 and MAPK9/JNK2 at early time points (Figure 5B), and it would be surprising (but not impossible) for this small increase to result in the predicted barrier-strengthening activity. In addition, there was a rise in MAPK8/JNK1 and MAPK9/JNK2 activation marks at late time points (180 and 240 minutes after thrombin treatment) in +thrombin condition (Figure 5B), despite the lack of predicted barrier activity. One possibility is this late JNK1/JNK2 activation mark may be related to a non-barrier function of JNK kinases. Alternatively, this may be a false-negative tKiR prediction. Overall, in most cases, increasing phosphorylation levels were observed immediately preceding or overlapping with the corresponding time windows of tKiR-predicted barrier activity for the ERK and the JNK pathways.

As noted above, MAPKAPK2/MK2 is a switch kinase, which has early barrier-weakening and late barrier-strengthening activity with thrombin treatment, but the late barrier-strengthening activity was disconnected after TNF pre-conditioning (Figure 4). As MAPKAPK2/MK2 can be activated by ERK, JNK, and p38 pathways, we used western blots to investigate how the three MAPKs signaling pathways may regulate this complex barrier phenotype. Consistent with the tKiR model (Figure 4), MAPKAPK2/MK2 activation increased at both early and late time points in +thrombin condition, but the late MAPKAPK2/MK2 activation was substantially reduced with TNF pre-conditioning (Figure 5B). The early MAPKAPK2/MK2 activation correlated with a rise in MAPK1/ERK2 and MAPK3/ERK1 activation (Figure 5A) and the late MAPKAPK2/MK2 activation correlated with a rise in MAPK8/JNK1 and MAPK9/JNK2 activation (Figure 5B). Moreover, the late MAPK8/9 (JNK1/2) activation marks were substantially reduced after TNF pre-conditioning (Figure 5B). This analysis indicates that the ERK pathway is more likely to contribute to early MAPKAPK2/MK2 activation, while the JNK pathway is more likely to contribute to late MAPKAPK2/MK2 activation.

### 3.5 Differential p38-driven activation of the MAPKAPK2/MK2 switch kinase in the two inflammatory conditions

To further investigate the MAPKAPK2/MK2 switch kinase, we examined the p38 pathway. The tKiR methodology predicted that a MAP3K20/ZAK-MAPK14/p38α-MAPKAPK2/MK2 pathway was involved in late barrier-strengthening in +thrombin condition, but it is disconnected with TNF pre-conditioning (Figure 4, Figure 6A). To test the model predictions, we probed the phosphorylation levels of MAP3K20/ZAK and the p38 isoforms by western blot. Control western blots showed that the total levels of p38 isoforms did not change in this period (Supplementary Figure 2A). The *MAP3K20*/*ZAK* gene is alternatively spliced into large (∼100 kDa) and small (∼55 kDa) isoforms (Gotoh et al., 2001). In HBMECs, phosphorylated MAP3K20/ZAK ran as two large isoforms (labeled isoforms 1 and 2) and a smaller isoform (labeled isoform 3) (Figure 6B). Whereas the smaller MAP3K20/ZAK isoform 3 (∼70 kDa) was activated at early time points (5 and 10 minutes after thrombin treatment), the larger MAP3K20/ZAK isoform 2 was activated at late time points (180 and 240 minutes after thrombin treatment), coincident with a period of predicted MAP3K20/ZAK barrier-strengthening activity in both inflammatory conditions (Figure 6B). Likewise, MAPK14/p38α was activated at both early and late time points in +thrombin condition, but the late activation was dampened with TNF pre-conditioning (Figure 6B). Taken together, these findings suggest the smaller MAP3K20/ZAK isoform 3 is likely responsible for the early p38 activation, and the larger MAP3K20/ZAK isoform 2 for the late barrier-strengthening functionality of MAPK14/p38α-MAPKAPK2/MK2. The dampening of MAPK14/p38α and MAPKAPK2/MK2 activation in the late barrier recovery phase upon TNF pre-conditioning is consistent with the model predictions that this late-stage barrier-strengthening function is rewired to other p38 isoforms that are unable to activate MAPKAPK2/MK2 when cells are pre-treated with TNF. Consequently, there is a dampening of both late JNK (Fig. 5B) and p38α activation after TNF pre-conditioning, which could contribute to the reprogramming of the MAPKAPK2/MK2 switch kinase.

**FIGURE 6.**
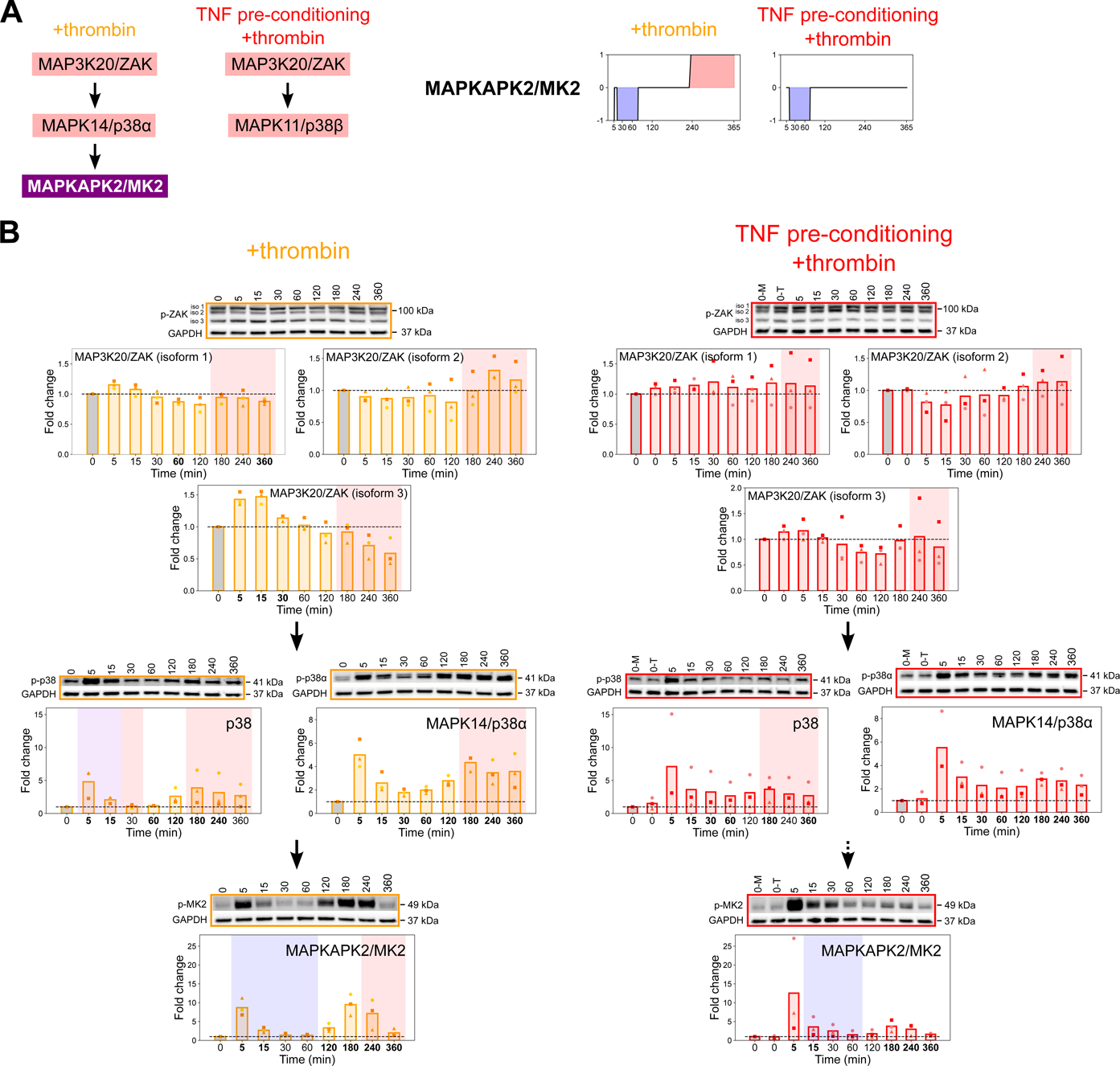
Differential p38-driven activation of the MAPKAPK2/MK2 switch kinase. HBMECs were stimulated with TNF (10 ng/ml) for 21 hours or left unstimulated, and cells were then treated with thrombin (5 nM) for the indicated times. **(A)** Left: tKiR model of how TNF pre-conditioning rewires the late barrier activity of the p38 pathway. Right: temporal barrier activity of MAPKAPK2/MK2 as predicted by tKiR. **(B)** Phosphorylation of members in the p38 pathway was probed by western blot in three biological replicates. Cells pre-conditioned with TNF but not treated with thrombin are labeled “0-T”. Data was normalized to non-treated, media only condition (“0-M”, gray bars). GAPDH was used as loading control. Symbols represent the fold change of individual biological replicates, and bars represent the mean fold change of three biological replicates. Blue and red shadings represent respectively barrier-weakening and barrier-strengthening activity, as predicted by tKiR, and purple shadings represent instances where different p38 kinases with barrier-weakening or barrier-strengthening functionality were predicted at that time point. The bolded timepoints indicate that the phosphorylation is different from the basal level (p-value from student’s t-test below 0.05 or all biological replicates reporting a fold change increasing by at least 20% compared to non-treated cells). See also Supplementary Figure 2 and Supplementary Table 3.

### 3.6 Construction of phosphosignaling networks that regulate barrier function

Among the kinases predicted by tKiR, there are many non-MAPK kinases (Supplementary Table 1). However, much less is known about non-MAPK signaling cascades and many kinases have been rarely studied. To discover new signaling pathways and crosstalk between pathways, we reasoned that kinases that are temporally related in barrier activity are more likely to act within a connected signaling module. Thus, we utilized self-organizing maps (SOMs) to systematically group kinases according to their temporal barrier activity. SOM is a dimensionality reduction method that can capture the topographic relationships of time series data (Kohonen, 1982). Kinases of similar temporal functionality were clustered into one group (called a “neuron”), and neurons having similar mean temporal behavior are more closely located on the SOM (Figure 2 (bottom), Figure 7A, Supplementary Figure 3A). To generate the SOMs, we used a grid size of 6×6 (resulting in up to 36 neurons). In +thrombin condition, kinases were grouped into 30 neurons and each neuron contained between 1 and 11 kinases (Figure 7A, Supplementary Figure 3B, Supplementary Table 2). In TNF pre-conditioning +thrombin condition, kinases were grouped into 35 neurons and each neuron contained between 1 and 9 kinases (Figure 7A, Supplementary Figure 3B, Supplementary Table 2).

**FIGURE 7.**
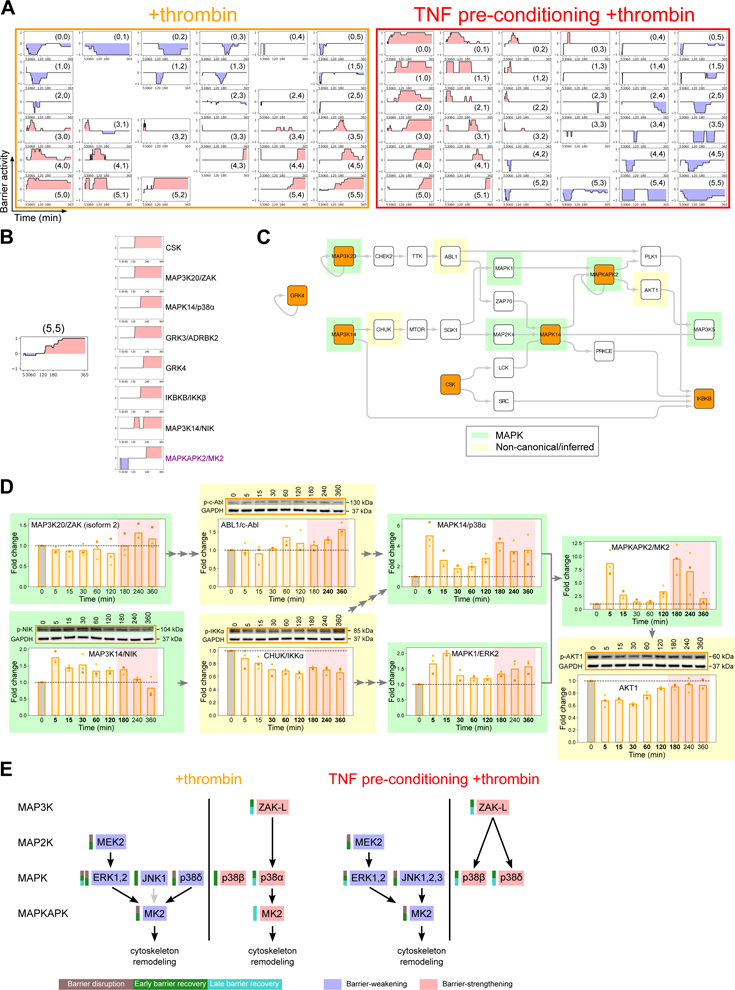
TREKING maps alternate routes to activation of the MAPKAPK2/MK2 switch kinase. **(A)** SOM clustering of tKiR predicted kinases with similar temporal functionality in +thrombin (left) or TNF pre-conditioning +thrombin (right) condition. Blue and red shadings represent barrier-weakening and barrier-strengthening functionality, respectively. In each neuron, the black line is the mean functionality of all the kinases within that neuron, with “+1” indicating all kinases are barrier-strengthening and “-1” indicating all kinases are barrier-weakening. Less than “1” means not all kinases in the neuron were predicted to be active in that time window. **(B)** A representative late barrier-strengthening-dominant neuron (neuron (5,5) in +thrombin condition) containing barrier-strengthening kinases of similar temporal barrier kinetics, as well as switch kinase MAPKAPK2/MK2. **(C)** Local phosphosignaling network reconstructed from neuron (5,5) in +thrombin condition. Kinases predicted by tKiR are labeled in orange; TREKING-inferred upstream/downstream kinases are unfilled. MAPKs and non-canonical/inferred kinases are highlighted in green and yellow, respectively. **(D)** Phosphorylation of tKiR-predicted and TREKING-inferred kinases within the network in **(C)** was probed by western blot in three biological replicates. Data was normalized to non-treated, media only condition (“0”, gray bars). Symbols represent the fold change of individual biological replicates, and bars represent the mean fold change of three biological replicates. Red shadings represent the late-stage (180-360 minutes after thrombin treatment) barrier-strengthening activity of neuron (5,5) as predicted by TREKING. Arrows represent the kinase connections in the network. The bolded timepoints indicate that the phosphorylation is different from the basal level (p-value from student’s t-test below 0.05 or all biological replicates reporting a fold change increasing by at least 20% compared to non-treated cells). **(E)** Cartoon model summarizing kinase regulation of the MAPKAPK2/MK2 switch kinase in the two inflammatory conditions. “ZAK-L” stands for the larger isoform (isoform 2) of MAP3K20/ZAK. See also Supplementary Figure 3, Supplementary Tables 2 and 3.

To investigate systems-level interconnections, we built TREKING models of barrier phosphosignaling networks (Figure 2, bottom). Specifically, for each neuron a local phosphosignaling network was built to describe the paths through which phosphosignals may propagate, using the kinase-substrate phosphorylation database PhosphoSitePlus^®^ (Hornbeck et al., 2015) to predict upstream and downstream kinases and to infer intermediate kinases in the pathways (Supplementary Figure 3C). We were able to build 26 local networks from the 30 neurons generated in +thrombin condition, and 27 local networks from the 35 neurons generated in TNF pre-conditioning +thrombin condition (Supplementary Table 2). The remaining neurons included only a single kinase that does not self-phosphorylate or multiple unconnected kinases. The size and topology of the phosphosignaling networks varied across neurons. In +thrombin condition, most networks had a maximum node-to-node shortest path length below 10 kinases, while the deepest networks built from neurons (0,0) and (3,3) had a maximum shortest path length of 14 kinases (Supplementary Figure 3C). In TNF pre-conditioning +thrombin condition, most networks had a maximum shortest path length between 3 and 9 kinases, except for the network built from neuron (2,4) that had a maximum shortest path length of 16 kinases (Supplementary Figure 3C).

### 3.7 TREKING maps alternate routes to activation of the MAPKAPK2/MK2 switch kinase

We used the TREKING models to further investigate the regulation of the MAPKAPK2/MK2 switch kinase. In the +thrombin condition, MAPKAPK2/MK2 clustered within a mid-to-late barrier-strengthening neuron (5,5). This neuron contained the MAP3K20/ZAK, MAPK14/p38α, and MAPKAPK2/MK2 components in the p38 pathway, as well as G-protein coupled receptors 3 and 4 (GRK3 and GRK4), C-terminal Src kinase (CSK; a negative regulator of Src-family kinases), and two members of non-canonical NF-kB signaling pathways (MAP3K14/NIK and inhibitor of nuclear factor kappa B kinase subunit beta (IKBKB/IKKβ)) (Figure 7B). In addition to the canonical MAP3K20/ZAK-MAPK14/p38α-MAPKAPK2/MK2 pathway (Figures 4 and 6), the TREKING model predicted several alternative pathways for late MAPKAPK2/MK2 activation (Figure 7C). One alternative pathway was predicted to link MAP3K20/ZAK and MAPK14/p38α via checkpoint kinase 2 (CHEK2), TTK protein kinase (TTK), ABL proto-oncogene 1 non-receptor tyrosine kinase (ABL1/c-Abl) and zeta chain of T cell receptor associated protein kinase 70 (ZAP70), instead of via the canonical MAPK pathway topology (MAP3K-MAP2K-MAPK) (Figure 7C). Other routes included a CSK-LCK pathway and a non-canonical NF-kB signaling pathway involving MAP3K14/NIK and inhibitor of nuclear factor kappa B kinase subunit alpha (CHUK/IKKα) as well as MAPK1/ERK2 (Figure 7C). Consistent with TREKING model predictions, activation marks increased in MAPK1/ERK2, ABL1/c-Abl, and CHUK/IKKα by western blot in the late barrier recovery phase (Figure 7D). One substrate of MAPKAPK2/MK2 is the heat shock protein HSP27 (HSPB1), which regulates actin microfilaments (Rada et al., 2021, Ronkina and Gaestel, 2022, Ronkina et al., 2008). The TREKING model also predicted that activated MAPKAPK2/MK2 leads to activation of MAP3K5/ASK1, suggesting a potential positive feedback loop, via AKT serine/threonine kinase 1 (AKT1), which exhibited increased phosphorylation in the late barrier recovery phase (Figure 7C-D).

MAPKAPK2/MK2 was no longer predicted to be a switch kinase with TNF pre-conditioning and clustered with an early barrier-weakening neuron (5,3) that also included MAPK3/ERK1, MKNK1, MAPK8/JNK1, p21 (Rac1) activated kinase 3 (PAK3), and Janus Kinase 1 (JAK1) (Supplementary Figure 4A). To better understand the early barrier-weakening activity of MAPKAPK2/MK2, we built a composite phosphosignaling network from neurons (5,3) and (5,4) (Supplementary Figure 4B). This TREKING model showed that ERK- and JNK-associated signaling may contribute to the activation of MAPKAPK2/MK2 in the barrier disruption phase and revealed crosstalk between ERK and JNK signaling pathways (Supplementary Figure 4B). One of the barrier-strengthening p38 MAPKs, MAPK14/p38α, was also inferred by the TREKING model that describes barrier-disruptive signaling, highlighting the complexity of kinase functionality and kinase-mediated signaling during different phases of barrier perturbation. Consistent with model predictions, increased levels of activated kinases in ERK, JNK, and p38 pathways, as well as the non-canonical, inferred kinase ABL1/c-Abl, were detected within the first 15 minutes of thrombin treatment by western blot (Supplementary Figure 4C).

The dichotomy between the signaling networks that activate MAPKAPK2/MK2 at early and late time points highlights the power of TREKING to uncover specific molecular mechanisms that mediate barrier disruption and recovery in a condition-specific way. Our analysis indicates that different kinases contribute to early and late MAPKAPK2/MK2 barrier activity, and the late barrier-strengthening pathway is rewired with TNF pre-conditioning. In +thrombin condition, both JNK and ERK signaling likely contribute to early MAPKAPK2/MK2 activation, while MAP3K20/ZAK-MAPK14/p38α signaling contribute to a second wave of MAPKAPK2/MK2 signaling and the resultant barrier-restorative activities (Figure 7E). In TNF pre-conditioning +thrombin condition, ERK and JNK signaling still both regulate early MAPKAPK2/MK2 activation, but with increased JNK activation (Figure 5). However, the late MAPK14/p38α-MAPKAPK2/MK2 barrier recovery pathway is reduced, and different p38 MAPK isoforms (MAPK11/p38β and MAPK13/p38δ) are predicted to play a more important role in barrier recovery in cells that are pre-conditioned with TNF (Figure 7E).

### 3.8 tKiR and TREKING expand the current understanding of kinases and associated phosphosignaling in barrier regulation

Most previous and ongoing research has focused on certain stages of barrier perturbation or sampled sparsely during a time course. We therefore asked how TREKING predictions compared to the existing literature in their breadth and temporal resolution. To do this, we visualized portions of the phosphosignaling networks that had been reported in the literature to regulate the endothelial barrier in response to thrombin (Supplementary Table 4). We then performed the same visualization with kinases that had been predicted by tKiR, and finally, by TREKING. Compared to the literature, TREKING substantially increased the number of kinases associated with thrombin-induced barrier regulation and was able to assemble most predicted kinases into time-resolved phosphosignaling networks (Figure 8, Supplementary File 1). This highlights the power of TREKING to broadly dissect functionally important phosphosignaling in barrier regulation, with high temporal resolution.

**FIGURE 8.**
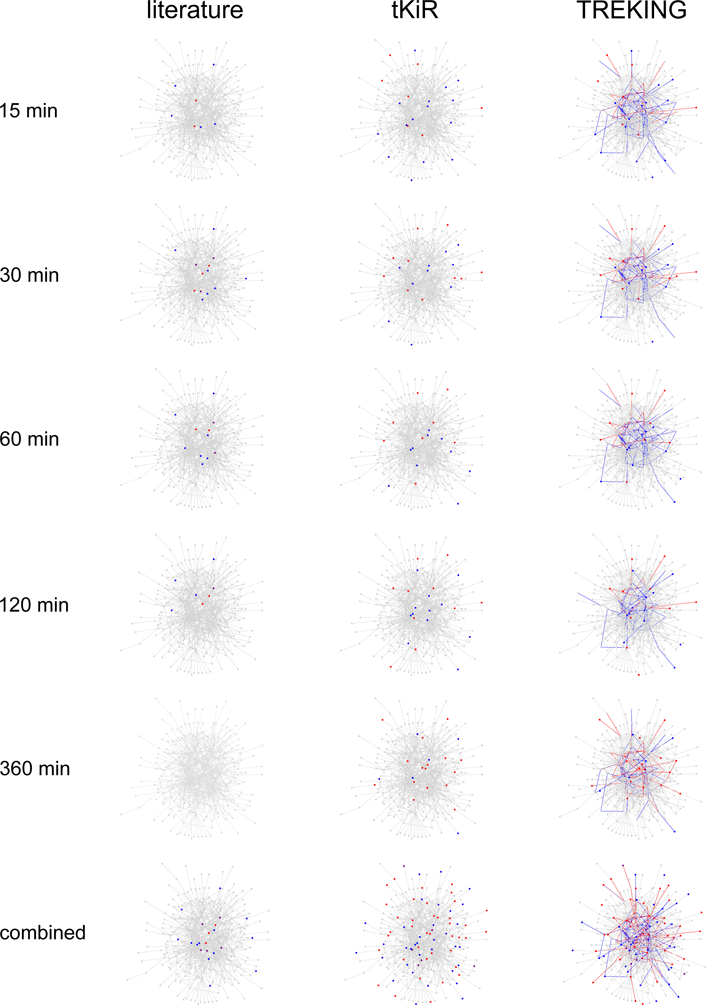
tKiR and TREKING expand the current understanding of kinases and associated phosphosignaling in barrier regulation. Comparison of literature-reported and model-predicted barrier-regulatory kinases and their connections at different stages of barrier perturbation. The background network in gray includes all kinase-kinase phosphorylation interactions from the kinase-substrate phosphorylation database PhosphoSitePlus^®^. Blue and red nodes are the kinases reported by literature or predicted by tKiR to have barrier-weakening and barrier-strengthening functionality, respectively; purple nodes are the kinases having conflicting literature reports. Blue and red edges are the interactions predicted by TREKING that are associated with only barrier-weakening and barrier-strengthening activity, respectively; purple edges are the interactions associated with both barrier-weakening and barrier-strengthening activities. The “combined” panel in the bottom is the composite literature reports or model predictions across all time points. The reported time points in the literature were expanded to 6-minute time windows (3 minutes before/after the reported time points) to account for any experimental variations. See also Supplementary Table 4 and Supplementary File 1.

## 4 Discussion

Kinase signaling pathways are highly implicated in endothelial barrier regulation (Mehta and Malik, 2006, Komarova et al., 2017, Kuppers et al., 2014) and consequently, kinase inhibitors are being explored for treatment of vascular injury (Aman et al., 2012, Botros et al., 2020, Rizzo et al., 2015). However, the experimental tools for reconstructing the complex signaling pathways in cells are underdeveloped, which has hindered therapeutic applications. Here, we introduce TREKING, which uses a small 28-panel kinase inhibitor screen, coupled with dynamic xCELLigence-based barrier measurements to broadly interrogate kinase signaling pathways involved in barrier regulation with temporal resolution. TREKING also allows us to study the interconnectedness in signaling networks that control the barrier.

In agreement with previous work (Mehta and Malik, 2006), our results using tKiR implicated multiple MAPK signaling pathways in regulating thrombin-induced barrier changes. Moreover, it was possible to assign barrier functions to specific MAPK pathways and in some instances to discern signaling cascades propagating specific barrier phenotypes. Our analysis indicates that the ERK and the JNK pathways were involved in the initial barrier disruption phase and the p38 pathway in mid-to-late acting barrier restoration. In each case, multiple kinases within each pathway were predicted and the barrier phenotypes followed the expected flow of phosphosignaling in the linear cascade.

Furthermore, TREKING accurately predicted both early and late barrier activities of the ERK and the p38 pathways, as validated by kinase activation by western blots, and it provided molecular insights into how late barrier activity in the ERK and the p38 cascades were rewired with TNF pre-conditioning. Beyond MAPKs, our findings highlight the complexity of endothelial barrier regulation with over 150 kinases being predicted to regulate thrombin-induced changes in barrier integrity.

Investigating the MAPKAPK2/MK2 switch kinase provides a useful example of the utility of TREKING to identify and dissect complex signaling pathways regulating barrier phenotypes after different inflammatory stimuli. The stress-activated MAPKAPK2/MK2 is involved in cytoskeleton remodeling associated with barrier permeability stimuli (Ronkina et al., 2008). Activation of MAPKAPK2/MK2 leads to phosphorylation of LIM domain kinase 1 (LIMK1) and HSPB1/HSP27, both of which regulate actin filament dynamics in endothelial cells (Rada et al., 2021, Gorovoy et al., 2005). Our analysis suggests that early barrier-weakening activity of MAPKAPK2/MK2 was controlled by the ERK and the JNK pathways and its late barrier-strengthening activity was regulated by a MAP3K20/ZAK-MAPK14/p38α signaling pathway, and crosstalk between p38, ERK, and JNK pathways. Consistent with the model predictions, both MAPK14/p38α and MAPKAPK2/MK2 activation were lessened at later time points when TNF pre-conditioning had taken place. Notably, the small and large MAP3K20/ZAK isoforms are both activated by stress stimuli and have been implicated in p38-mediated regulation of actin stress fibers via different mechanisms (Gotoh et al., 2001, Nordgaard et al., 2022). The small and large MAP3K20/ZAK isoforms are phosphorylated with different kinetics after thrombin stimulation in HBMECs, suggesting another level of regulatory control. Together, this analysis suggests a complex regulation of MAPKAPK2/MK2 that is influenced by several MAPK pathways during both the barrier disruption and repair phases and is partially rewired by TNF pre-conditioning. Crosstalk between MAPK pathways in mediating barrier permeability has been implicated previously, but lack of molecular details describing their interconnections, associated temporal kinetics, and the role of non-canonical players, make it challenging to fully understand their regulatory mechanisms in the context of barrier function (Paumelle et al., 2000, Zhang et al., 2001, Surapisitchat et al., 2001).

Several examples illustrate additional molecular and kinetic details that can be obtained from TREKING in contrast to traditional, less systematic approaches. These details include shifts in the timing of core MAPK signaling pathways (ERK and JNK pathways), time-resolved barrier phenotypes, condition-specific phosphosignaling networks (the MAPKAPK2/MK2 switch kinase, which we investigated in detail is one of several predicted examples), and engagement of non-canonical kinase signaling pathways. TREKING also highlights how the ERK, JNK, and p38 MAPK pathways are altered in TNF pre-conditioned cells. These differences are often buried by conventional investigations that use tools such as genetic knockdown or knockout, which respond on the time scale of days rather than minutes. This highlights the importance of developing tools to systematically study the molecular details of kinase-driven signaling and to capture the differences in cell signaling between conditions.

Despite its power to elucidate kinase signaling with temporal resolution, TREKING is not without limitations. Firstly, the kinase-compound biochemical data used for tKiR contains information only on a subset of the human kinome (291of 518 kinases) (Anastassiadis et al., 2011). Moreover, some kinases are targeted less frequently than others by the 28-kinase inhibitor panel, and this skewing of the data can lead to both false positive and false negative predictions. To partially overcome this limitation, TREKING reconstructs phosphosignaling networks and infers kinases not predicted by tKiR and therefore can fill in missing gaps by leveraging kinase proteomic datasets. Most kinase signaling pathways are much less studied than the MAPKs and therefore TREKING is also a discovery tool to identify new pathways that will need to be validated in the future. As additional kinase-compound biochemical data is available, these models can be refined, or the panel of kinase inhibitors can be expanded to improve resolution on the understudied kinases or kinase gene families. Secondly, PhosphoSitePlus^®^, the knowledgebase used for building the phosphosignaling networks, despite being the most comprehensive catalog available to us, does not cover the complete kinase-substrate phosphorylation interactions in human cells, nor is it specific to endothelial cells. As more advanced phosphoproteomics datasets and tools to dissect kinase-substrate relationships become available (Johnson et al., 2023), the subsequent iterations of TREKING will incorporate this comprehensive knowledge to further refine the models. Thirdly, information such as protein abundance, subcellular localization, translation, degradation, and dephosphorylation, were not incorporated in building TREKING models in the current study, due to the lack of extensive knowledge on kinase abundance and kinetics in brain endothelial cells. The TREKING models can be further refined as this information becomes available.

In conclusion, our results reveal commonalities and divergences of kinase signaling in response to classic pro-inflammatory stimuli, and insight into signaling pathways with barrier-weakening or barrier-strengthening activity and crosstalk between them. TREKING allows 1) assessment of time-resolved kinase functionality associated with barrier regulation, 2) construction of phenotype-driving kinase-mediated phosphosignaling networks that recapitulate signaling events at different stages of barrier perturbation, and 3) systematic comparison of functional phosphosignaling networks between conditions. The global, time-resolved and mechanistic details revealed by TREKING has ramifications beyond our fundamental understanding of phosphosignaling during inflammation.

Beyond providing a deeper understanding into control of the vasculature, TREKING is broadly generalizable to other cellular systems with kinetic datasets, and could be applied towards developing a molecular, kinetically resolved picture of other cellular phenotypes.

## Data Availability Statement

Data and code generated in this study are available in the main text or the supplementary material. Further information and requests for resources and reagents should be directed to and will be fulfilled by the corresponding author, Alexis Kaushansky.

## Author Contributions

Conceptualization, L.W., S.D., J.D.S., and A.K.; Funding acquisition, J.D.S. and A.K.; Investigation, L.W. and S.D.; Methodology, L.W., S.D., and K.V.; Supervision, J.D.S. and A.K.; Writing – Original Draft, L.W. with help from S.D., J.D.S. and A.K.; Writing – Review & Editing, L.W., S.D., J.D.S., and A.K.

## Acknowledgments

The authors would like to thank Mary-Margaret Dols for experimental guidance.

## References

Aman, J., Van Bezu, J., Damanafshan, A., Huveneers, S., Eringa, E. C., Vogel, S. M., Groeneveld, A. B., Vonk Noordegraaf, A., Van Hinsbergh, V. W. & Van Nieuw Amerongen, G. P. 2012. Effective treatment of edema and endothelial barrier dysfunction with imatinib. Circulation, 126, 2728–38.

Anastassiadis, T., Deacon, S. W., Devarajan, K., Ma, H. & Peterson, J. R. 2011. Comprehensive assay of kinase catalytic activity reveals features of kinase inhibitor selectivity. Nat Biotechnol, 29, 1039–45.

Anrather, D., Millan, M. T., Palmetshofer, A., Robson, S. C., Geczy, C., Ritchie, A. J., Bach, F. H. & Ewenstein, B. M. 1997. Thrombin activates nuclear factor-kappaB and potentiates endothelial cell activation by TNF. J Immunol, 159, 5620–8.

Arang, N., Kain, H. S., Glennon, E. K., Bello, T., Dudgeon, D. R., Walter, E. N. F., Gujral, T. S. & Kaushansky, A. 2017. Identifying host regulators and inhibitors of liver stage malaria infection using kinase activity profiles. Nat Commun, 8, 1232.

Beguin, E. P., van den Eshof, B. L., Hoogendijk, A. J., Nota, B., Mertens, K., Meijer, A. B. & van den Biggelaar, M. 2019. Integrated proteomic analysis of tumor necrosis factor alpha and interleukin 1beta-induced endothelial inflammation. J Proteomics, 192, 89–101.

Birukova, A. A., Tian, X., Tian, Y., Higginbotham, K. & Birukov, K. G. 2013. Rap-afadin axis in control of Rho signaling and endothelial barrier recovery. Mol Biol Cell, 24, 2678–88.

Botros, L., Pronk, M. C. A., Juschten, J., Liddle, J., Morsing, S. K. H., Van Buul, J. D., Bates, R. H., Tuinman, P. R., Van Bezu, J. S. M., Huveneers, S., Bogaard, H. J., Van Hinsbergh, V. W. M., Hordijk, P. L. & Aman, J. 2020. Bosutinib prevents vascular leakage by reducing focal adhesion turnover and reinforcing junctional integrity. J Cell Sci, 133.

Cargnello, M. & Roux, P. P. 2011. Activation and function of the MAPKs and their substrates, the MAPK-activated protein kinases. Microbiol Mol Biol Rev, 75, 50–83.

Casado, P., Rodriguez-Prados, J. C., Cosulich, S. C., Guichard, S., Vanhaesebroeck, B., Joel, S. & Cutillas, P. R. 2013. Kinase-substrate enrichment analysis provides insights into the heterogeneity of signaling pathway activation in leukemia cells. Sci Signal, 6, rs6.

Daneman, R. & Prat, A. 2015. The blood-brain barrier. Cold Spring Harb Perspect Biol, 7, a020412.

Dankwa, S., Dols, M. M., Wei, L., Glennon, E. K. K., Kain, H. S., Kaushansky, A. & Smith, J. D. 2021. Exploiting polypharmacology to dissect host kinases and kinase inhibitors that modulate endothelial barrier integrity. Cell Chem Biol.

Garcia, J. G., Liu, F., Verin, A. D., Birukova, A., Dechert, M. A., Gerthoffer, W. T., Bamberg, J. R. & English, D. 2001. Sphingosine 1-phosphate promotes endothelial cell barrier integrity by Edg-dependent cytoskeletal rearrangement. J Clin Invest, 108, 689–701.

Gorovoy, M., Niu, J., Bernard, O., Profirovic, J., Minshall, R., Neamu, R. & Voyno-Yasenetskaya, T. 2005. LIM kinase 1 coordinates microtubule stability and actin polymerization in human endothelial cells. J Biol Chem, 280, 26533–42.

Gotoh, I., Adachi, M. & Nishida, E. 2001. Identification and characterization of a novel MAP kinase kinase kinase, MLTK. J Biol Chem, 276, 4276–86.

Gujral, T. S., Peshkin, L. & Kirschner, M. W. 2014. Exploiting polypharmacology for drug target deconvolution. Proc Natl Acad Sci U S A, 111, 5048–53.

Han, J., Zhang, G., Welch, E. J., Liang, Y., Fu, J., Vogel, S. M., Lowell, C. A., Du, X., Cheresh, D. A., Malik, A. B. & Li, Z. 2013. A critical role for Lyn kinase in strengthening endothelial integrity and barrier function. Blood, 122, 4140–9.

Hornbeck, P. V., Zhang, B., Murray, B., Kornhauser, J. M., Latham, V. & Skrzypek, E. 2015. PhosphoSitePlus, 2014: mutations, PTMs and recalibrations. Nucleic Acids Res, 43, D512-20.

Johnson, J. L., Yaron, T. M., Huntsman, E. M., Kerelsky, A., Song, J., Regev, A., Lin, T. Y., Liberatore, K., Cizin, D. M., Cohen, B. M., Vasan, N., Ma, Y., Krismer, K., Robles, J. T., van de Kooij, B., Van Vlimmeren, A. E., Andree-Busch, N., Kaufer, N. F., Dorovkov, M. V., Ryazanov, A. G., Takagi, Y., Kastenhuber, E. R., Goncalves, M. D., Hopkins, B. D., Elemento, O., Taatjes, D. J., Maucuer, A., Yamashita, A., Degterev, A., Uduman, M., Lu, J., Landry, S. D., Zhang, B., Cossentino, I., Linding, R., Blenis, J., Hornbeck, P. V., Turk, B. E., Yaffe, M. B. & Cantley, L. C. 2023. An atlas of substrate specificities for the human serine/threonine kinome. Nature, 613, 759–766.

Klomp, J. E., Shaaya, M., Matsche, J., Rebiai, R., Aaron, J. S., Collins, K. B., Huyot, V., Gonzalez, A. M., Muller, W. A., Chew, T. L., Malik, A. B. & Karginov, A. V. 2019. Time-Variant SRC Kinase Activation Determines Endothelial Permeability Response. Cell Chem Biol, 26, 1081–1094 e6.

Knezevic, N., Tauseef, M., Thennes, T. & Mehta, D. 2009. The G protein betagamma subunit mediates reannealing of adherens junctions to reverse endothelial permeability increase by thrombin. J Exp Med, 206, 2761–77.

Kohonen, T. 1982. Self-organized formation of topologically correct feature maps. Biological cybernetics, 43, 59–69.

Komarova, Y. A., Kruse, K., Mehta, D. & Malik, A. B. 2017. Protein Interactions at Endothelial Junctions and Signaling Mechanisms Regulating Endothelial Permeability. Circ Res, 120, 179–206.

Kuppers, V., Vockel, M., Nottebaum, A. F. & Vestweber, D. 2014. Phosphatases and kinases as regulators of the endothelial barrier function. Cell Tissue Res, 355, 577–86.

Liu, Y., Pelekanakis, K. & Woolkalis, M. J. 2004. Thrombin and tumor necrosis factor alpha synergistically stimulate tissue factor expression in human endothelial cells: regulation through c-Fos and c-Jun. J Biol Chem, 279, 36142–7.

Marcos-Ramiro, B., Garcia-Weber, D. & Millan, J. 2014. TNF-induced endothelial barrier disruption: beyond actin and Rho. Thromb Haemost, 112, 1088–102.

Mcverry, B. J. & Garcia, J. G. 2004. Endothelial cell barrier regulation by sphingosine 1-phosphate. J Cell Biochem, 92, 1075–85.

Mehta, D. & Malik, A. B. 2006. Signaling mechanisms regulating endothelial permeability. Physiol Rev, 86, 279–367.

Miller, L. H., Ackerman, H. C., Su, X. Z. & Wellems, T. E. 2013. Malaria biology and disease pathogenesis: insights for new treatments. Nat Med, 19, 156–67.

Minami, T., Sugiyama, A., Wu, S. Q., Abid, R., Kodama, T. & Aird, W. C. 2004. Thrombin and phenotypic modulation of the endothelium. Arterioscler Thromb Vasc Biol, 24, 41–53.

Morrison, D. K. 2012. MAP kinase pathways. Cold Spring Harb Perspect Biol, 4.

Nordgaard, C., Vind, A. C., Stonadge, A., Kjobsted, R., Snieckute, G., Antas, P., Blasius, M., Reinert, M. S., del Val, A. M., Bekker-Jensen, D. B., Haahr, P., Miroshnikova, Y. A., Mazouzi, A., Falk, S., Perrier-Groult, E., Tiedje, C., Li, X., Jakobsen, J. R., Jorgensen, N. O., Wojtaszewski, J. F., Mallein-Gerin, F., Andersen, J. L., Pennisi, C. P., Clemmensen, C., Kassem, M., Jafari, A., Brummelkamp, T., Li, V. S., Wickstrom, S. A., Olsen, J. V., Blanco, G. & Bekker-Jensen, S. 2022. ZAKbeta is activated by cellular compression and mediates contraction-induced MAP kinase signaling in skeletal muscle. EMBO J, 41, e111650.

Oldenburg, J. & de Rooij, J. 2014. Mechanical control of the endothelial barrier. Cell Tissue Res, 355, 545–55.

Paumelle, R., Tulasne, D., Leroy, C., Coll, J., Vandenbunder, B. & Fafeur, V. 2000. Sequential activation of ERK and repression of JNK by scatter factor/hepatocyte growth factor in madin-darby canine kidney epithelial cells. Mol Biol Cell, 11, 3751–63.

Pimienta, G. & Pascual, J. 2007. Canonical and alternative MAPK signaling. Cell Cycle, 6, 2628–32.

Rada, C. C., Mejia-Pena, H., Grimsey, N. J., Canto Cordova, I., Olson, J., Wozniak, J. M., Gonzalez, D. J., Nizet, V. & Trejo, J. 2021. Heat shock protein 27 activity is linked to endothelial barrier recovery after proinflammatory GPCR-induced disruption. Sci Signal, 14, eabc1044.

Radeva, M. Y. & Waschke, J. 2018. Mind the gap: mechanisms regulating the endothelial barrier. Acta Physiol (Oxf), 222.

Rizzo, A. N., Aman, J., van Nieuw Amerongen, G. P. & Dudek, S. M. 2015. Targeting Abl kinases to regulate vascular leak during sepsis and acute respiratory distress syndrome. Arterioscler Thromb Vasc Biol, 35, 1071–9.

Ronkina, N. & Gaestel, M. 2022. MAPK-Activated Protein Kinases: Servant or Partner? Annu Rev Biochem, 91, 505–540.

Ronkina, N., Kotlyarov, A. & Gaestel, M. 2008. MK2 and MK3--a pair of isoenzymes? Front Biosci, 13, 5511–21.

Schafer, A., Gjerga, E., Welford, R. W., Renz, I., Lehembre, F., Groenen, P. M., Saez-Rodriguez, J., Aebersold, R. & Gstaiger, M. 2019. Elucidating essential kinases of endothelin signalling by logic modelling of phosphoproteomics data. Mol Syst Biol, 15, e8828.

Surapisitchat, J., Hoefen, R. J., Pi, X., Yoshizumi, M., Yan, C. & Berk, B. C. 2001. Fluid shear stress inhibits TNF-alpha activation of JNK but not ERK1/2 or p38 in human umbilical vein endothelial cells: Inhibitory crosstalk among MAPK family members. Proc Natl Acad Sci U S A, 98, 6476–81.

Tiruppathi, C., Naqvi, T., Sandoval, R., Mehta, D. & Malik, A. B. 2001. Synergistic effects of tumor necrosis factor-alpha and thrombin in increasing endothelial permeability. Am J Physiol Lung Cell Mol Physiol, 281, L958–68.

Vaga, S., Bernardo-Faura, M., Cokelaer, T., Maiolica, A., Barnes, C. A., Gillet, L. C., Hegemann, B., Van Drogen, F., Sharifian, H., Klipp, E., Peter, M., Saez-Rodriguez, J. & Aebersold, R. 2014. Phosphoproteomic analyses reveal novel cross-modulation mechanisms between two signaling pathways in yeast. Mol Syst Biol, 10, 767.

van den Biggelaar, M., Hernandez-Fernaud, J. R., van den Eshof, B. L., Neilson, L. J., Meijer, A. B., Mertens, K. & Zanivan, S. 2014. Quantitative phosphoproteomics unveils temporal dynamics of thrombin signaling in human endothelial cells. Blood, 123, e22–36.

Vouret-Craviari, V., Bourcier, C., Boulter, E. & van Obberghen-Schilling, E. 2002. Distinct signals via Rho GTPases and Src drive shape changes by thrombin and sphingosine-1-phosphate in endothelial cells. J Cell Sci, 115, 2475–84.

Zhang, H., Shi, X., Hampong, M., Blanis, L. & Pelech, S. 2001. Stress-induced inhibition of ERK1 and ERK2 by direct interaction with p38 MAP kinase. J Biol Chem, 276, 6905–8.

Zhao, Z., Nelson, A. R., Betsholtz, C. & Zlokovic, B. V. 2015. Establishment and Dysfunction of the Blood-Brain Barrier. Cell, 163, 1064–1078.

Zhu, L., Yu, Q., Gao, P., Liu, Q., Luo, X., Jiang, G., Ji, R., Yang, R., Ma, X., Xu, J., Yuan, H., Zhou, J. & An, H. 2020. TAOK1 positively regulates TLR4-induced inflammatory responses by promoting ERK1/2 activation in macrophages. Mol Immunol, 122, 124–131.

