## Supplementary Material for "Interrogating endothelial barrier regulation by temporally resolved kinase network generation"

**Supplementary Figure 1.** **xCELLigence data on HBMECs treated with thrombin (with or without TNF pre-conditioning), related to Figure 1.** **(A)** Cell index data from Figure 1A plotted with no normalization. Left: time course data from the addition of TNF. Right: time course data from the addition of thrombin and subsequent 6 hours. **(B)** Cell index data from Figure 1A plotted with timepoint normalization and no baseline normalization. Left: normalized time course data from the addition of TNF. Right: normalized time course data from the addition of thrombin and subsequent 6 hours. **(C)** DMSO vehicle used for kinase inhibitors has minimal effect on permeability of HBMECs treated with thrombin (with or without TNF pre-conditioning). Left: cell index data plotted with no normalization. Right: cell index data plotted with timepoint normalization and no baseline normalization.

**Supplementary Figure 2. Total protein levels of ERK, JNK and p38 kinases are stable across the time course, related to Figures 5, 6 and 7.** **(A)** Total protein levels of ERK, JNK and p38 kinases were probed by western blot in three biological replicates across the 6-hour time course after thrombin treatment (with or without TNF pre-conditioning). Cells pre-conditioned with TNF but not treated with thrombin are labeled “0-T”. Data was normalized to non-treated, media only condition (“0-M”, gray bars). GAPDH was used as loading control. **(B)** Phosphorylation of ERK, JNK and p38 kinases was quantified by first normalizing to their total protein levels and then normalizing to non-treated, media only condition (“0-M”, gray bars) for fold change from the basal level. The fold changes are similar to quantification by normalizing to GAPDH (Figure 5, Figure 6B, Figure 7D). Symbols represent the fold change of individual biological replicates, and bars represent the mean fold change of three biological replicates. The bolded timepoints indicate that the phosphorylation is different from the basal level. See also Supplementary Table 3.

**Supplementary Figure 3. Building of local phosphosignaling networks based on kinase functional kinetics, related to Figure 7.** **(A)** Distance maps of SOM clustering in +thrombin (left) and TNF pre-conditioning +thrombin (right) conditions. Each cell in the grid represents the normalized sum of the Euclidean distances between a neuron and its neighboring neurons. **(B)** Distribution of the number of kinases assigned to the SOM neurons. **(C)** Left: example of how a local phosphosignaling network was built for each neuron by searching for the shortest paths between any pair of kinases within that neuron using the kinase-substrate phosphorylation database on PhosphoSitePlus^®^ as the background network. Right: the maximum path length in the local phosphosignaling networks. In panels **(B)** and **(C)**, the individual neurons are listed within the respective bar graphs.

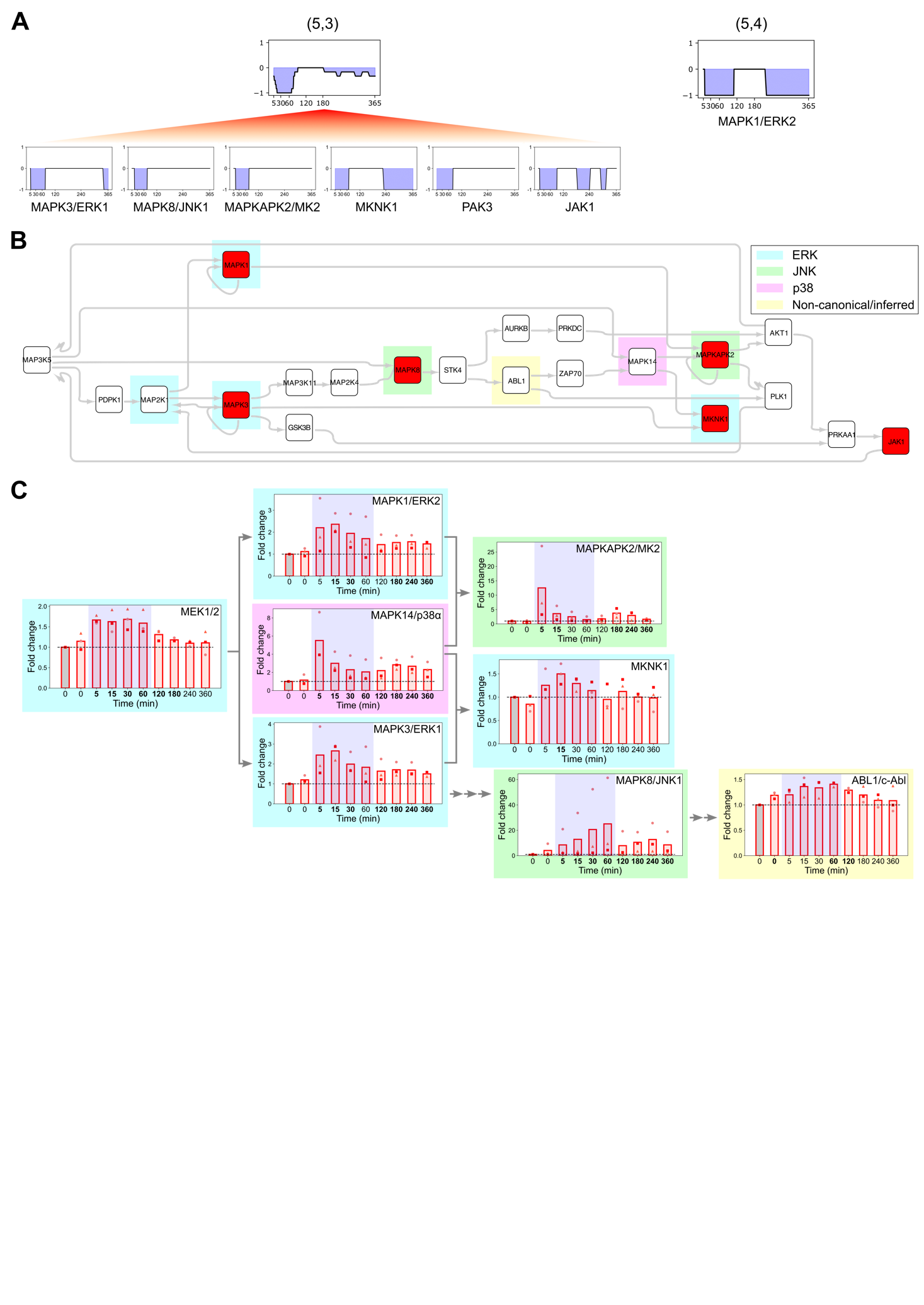

**Supplementary Figure 4. TREKING predicts kinase-mediated signaling pathways functionally important for barrier disruption, related to Figure 7.** **(A)** Representative barrier-weakening neurons (5,3) and (5,4) generated in TNF pre-conditioning +thrombin condition containing barrier-weakening kinases of similar temporal barrier kinetics. **(B)** Composite phosphosignaling network reconstructed from neurons (5,3) and (5,4) in TNF pre-conditioning +thrombin condition. Kinases predicted by tKiR are labeled in red; TREKING-inferred kinases are unfilled. Kinases associated with ERK, JNK and p38 signaling are highlighted in cyan, green and purple, respectively, and non-canonical/inferred kinases are highlighted in yellow. **(C)** Phosphorylation of tKiR-predicted and TREKING-inferred kinases within the network was probed by western blot (5-360 minutes after thrombin treatment) in three biological replicates. Data was normalized to non-treated, media only condition (“0”, gray bars). Symbols represent the fold change of individual biological replicates, and bars represent the mean fold change of three biological replicates. Blue shadings represent the earlier stage (5-60 minutes after thrombin treatment) of barrier-weakening activity as predicted by TREKING. Arrows represent the kinase connections in the network. The bolded timepoints indicate that the phosphorylation is different from the basal level.

**Supplementary Table 1. Predicted temporal kinase functionality in +thrombin and TNF pre-conditioning +thrombin conditions, related to Figure 3.** This is a Microsoft Excel workbook containing two spreadsheets. The column headers represent the midpoint of each 5-minute time window where tKiR was performed. The binarized numbers stand for the tKiR predicted barrier function (“-1”: barrier-weakening; “1”: barrier-strengthening; “0”: not predicted by tKiR).

**Supplementary Table 2. SOM assignment of tKiR-predicted kinases and edges of phosphosignaling networks built in +thrombin and TNF pre-conditioning +thrombin conditions, related to Figure 7.** This is a Microsoft Excel workbook containing four spreadsheets.

**Supplementary Table 3. Antibody information and western blot results on the kinases validated in this study, related to Figures 5, 6 and 7.** This is a Microsoft Excel workbook containing 17 spreadsheets of the antibody information and densitometry results for the 15 kinase antibody targets.

**Supplementary Table 4. Literature search on protein kinases reported to regulate endothelial barrier integrity in response to thrombin, related to Figure 8.** PubMed ID of the article is followed by the barrier functionality reported in the corresponding article (“-1”: barrier-weakening; “1”: barrier-strengthening). This is a Microsoft Excel workbook containing one spreadsheet.

**Supplementary File 1. Movie comparing literature-reported with tKiR- and TREKING-predicted kinases and signaling pathways important for barrier regulation.** This is an MPEG-4 movie file.

**Supplementary File 2. Jupyter notebook containing the scripts for tKiR, SOM generation and network construction.** Also included is the script for making frames for the supplementary movie (Supplementary File 1).
