## Supplementary File 2 for "Interrogating endothelial barrier regulation by temporally resolved kinase network generation": Supplemental file 2.ipynb

```
# %% [code]
import os
import glob
import numpy as np
import pandas
import scipy
import random
import csv
from minisom import MiniSom
import networkx as nx
import glmnet_python
from glmnet import glmnet 
from glmnetCoef import glmnetCoef
from glmnetPredict import glmnetPredict
import matplotlib
import matplotlib.pyplot as plt
from matplotlib.gridspec import GridSpec

# %% [markdown]
# Temporal KiR

# %% [code]
path = "..."

ResidualActivity = pandas.read_csv(path + "ResidualActivity.txt", sep='\t')
print(ResidualActivity.shape)

data = pandas.read_csv(path + "KinaseInhibitorScreens_AUC.txt", sep='\t')
print(data.shape)

drugs = data['Drug']
drug_list = drugs.unique()

x = ResidualActivity.iloc[:, 1:].to_numpy()
Y = data.iloc[:, 1:].to_numpy()

alpha = 0.8
nlambda = 1e3

coeffs = np.zeros((x.shape[1] + 1, Y.shape[1]), dtype=float)

for t in range(Y.shape[1]):
    
    y = Y[:, t]

    fit = glmnet(x = x.copy(), y = y.copy(), family = 'gaussian', alpha = alpha, nlambda = nlambda)

    lambdas = np.array(fit['lambdau'])

    mse = np.zeros((len(lambdas), len(drug_list)))
    mae = np.zeros((len(lambdas), len(drug_list)))

    for index, i_lambda in enumerate(lambdas):
        i = 0
        for drug in drug_list:

            x_validate = x.drop(drugs[drugs != drug].index).to_numpy()
            y_true = y.drop(drugs[drugs != drug].index).to_numpy()

            y_predict = glmnetPredict(fit, x_validate, ptype = 'response', s = scipy.float64([i_lambda]))

            mse[index, i] = np.sum(np.square(np.subtract(y_predict, y_true)))
            mae[index, i] = np.sum(abs(np.subtract(y_predict, y_true)))

            i += 1

    lambda_min = lambdas[np.argmin(np.mean(mse, axis=1))]
    # print(np.argmin(np.mean(mse, axis=1)), lambda_min)

    coef = glmnetCoef(fit, s = scipy.float64([lambda_min]), exact = False)
    # print('Non-zero coefficients: ', np.count_nonzero(coef[1:]))
    
    for i in range(len(coef)):
        coeffs[i, t] = coef[i]

np.savetxt("TemporalKiR_AUC.txt", coeffs, delimiter='\t')


# %% [markdown]
# Self-organizing map

# %% [code]
# Train

path = "..."

data = pandas.read_csv(path + "TemporalKiR_AUC.binary.txt", sep='\t')

coeffs = pandas.DataFrame(data.iloc[:, 1:])
coeffs = coeffs.to_numpy()
print(coeffs.shape)

(x_dim, y_dim) = (6, 6)

som = MiniSom(x_dim, y_dim, coeffs.shape[1], sigma=1.0, learning_rate=0.2, neighborhood_function='gaussian', random_seed=1)
som.pca_weights_init(coeffs)
som.train_batch(coeffs, num_iteration=10000, verbose=True)


# %% [code]
# Distance map

som_labels = np.empty(coeffs.shape, object)

for i in range(coeffs.shape[0]):
    for j in range(coeffs.shape[1]):
        som_labels[i, j] = som.winner(coeffs[i, j])

matplotlib.style.use('default')
plt.rcParams["font.family"] = "Arial"

plt.imshow(som.distance_map(), aspect='auto', cmap='RdYlBu')
plt.colorbar(fraction=0.05)
plt.grid(False)

plt.savefig(path + "som(6,6).DistanceMap.pdf", format='pdf')


# %% [code]
# Generate SOMs

win_map = som.win_map(coeffs)

with open(path + "NeuronAssignment.txt", 'w') as f:

    writer = csv.writer(f, delimiter='\t')
    for k, v in win_map.items():
        writer.writerow([k] + v)

matplotlib.style.use('default')
plt.figure(figsize=(16, 12))

plt.rcParams["font.family"] = "Arial"
the_grid = GridSpec(x_dim, y_dim)
ticks = [5, 30, 60, 120, 180, 240, 360]

time = data.iloc[:, 1:].columns.to_numpy(dtype=float)

for position in win_map.keys():
    plt.subplot(the_grid[position[0], position[1]])
    
    plt.plot(time, np.min(win_map[position], axis=0), color='gray', alpha=0.5)
    plt.plot(time, np.mean(win_map[position], axis=0), color='black')
    plt.plot(time, np.max(win_map[position], axis=0), color='gray', alpha=0.5)
    
    plt.xticks(ticks=ticks, labels=ticks)
    plt.ylim(-1.1, 1.1)
    
    plt.fill_between(time, np.mean(win_map[position], axis=0), where=(np.mean(win_map[position], axis=0)<0.0), color='blue', alpha=0.30, interpolate=True)
    plt.fill_between(time, np.mean(win_map[position], axis=0), where=(np.mean(win_map[position], axis=0)>0.0), color='red', alpha=0.30, interpolate=True)
    plt.grid(False)

plt.savefig(path + "som(6,6).pdf", format='pdf')


# %% [markdown]
# Path search

# %% [code]
path = "..."

KSIN = pandas.read_csv(path + "PhosphoSitePlus_KSdata.txt", sep='\t')
KSIN = pandas.DataFrame(KSIN, columns=['Kinase Gene ID', 'Kinase Uniprot ID', 'Substrate Gene ID', 'Substrate Uniprot ID'])
print(KSIN.shape)

selfP = pandas.read_csv(path + "PhosphoSitePlus_selfP.txt", sep='\t')
selfP = pandas.DataFrame(selfP, columns=['Gene ID', 'Uniprot ID'])
print(selfP.shape)

all_files = glob.glob(path + "neurons/")

for filename in all_files:

    nodes = pandas.read_csv(filename, sep='\t')
    nodes = pandas.DataFrame(nodes, columns=['Gene ID'])

    G_KS = nx.from_pandas_edgelist(KSIN, source='Kinase Gene ID', target='Substrate Gene ID', edge_attr=None, create_using=nx.DiGraph)

    KS_path = pandas.DataFrame(columns=['Kinase Gene ID', 'Substrate Gene ID', 'Path Length', 'Path'])
    for i in range(nodes.shape[0]):
        for j in range(nodes.shape[0]):
            if i == j and nodes['Gene ID'].iloc[i] in selfP['Gene ID'].values:
                KS_path = KS_path.append(pandas.DataFrame({'Kinase Gene ID': [nodes['Gene ID'].iloc[i]],  
                                                        'Substrate Gene ID': [nodes['Gene ID'].iloc[j]],
                                                        'Path Length': [0],
                                                        'Path': [nodes['Gene ID'].iloc[i]]}), ignore_index=True)
            else:
                if nodes['Gene ID'].iloc[i] in KSIN.values and nodes['Gene ID'].iloc[j] in KSIN.values:
                    try:
                        path_length = nx.shortest_path_length(G_KS, source=nodes['Gene ID'].iloc[i], target=nodes['Gene ID'].iloc[j])
                        path = nx.shortest_path(G_KS, source=nodes['Gene ID'].iloc[i], target=nodes['Gene ID'].iloc[j])
                    except:
                        pass
                    else:
                        KS_path = KS_path.append(pandas.DataFrame({'Kinase Gene ID': [nodes['Gene ID'].iloc[i]],  
                                                                'Substrate Gene ID': [nodes['Gene ID'].iloc[j]],
                                                                'Path Length': [path_length],
                                                                'Path': [path]}), ignore_index=True)

    KS_path = KS_path.sort_values(by=['Path Length'], axis=0, ascending=True)
            
    DataFrame.to_csv(KS_path, filename + ".paths.txt", index=False, sep='\t', na_rep='')


# %% [markdown]
# Compare between literature, temporal KiR and TREKING

# %% [code]
# Generate individual frames

path = "..."

KSIN = pandas.read_csv(path + "PhosphoSitePlus_KSdata.txt", sep='\t')
KSIN = pandas.DataFrame(KSIN, columns=['Kinase Gene ID', 'Substrate Gene ID', 'Edge'])
print(KSIN.shape)

activity_literature = pandas.read_csv(path + "literature.txt", sep='\t', header=None)
activity_KiR = pandas.read_csv(path + "KiR.txt", sep='\t', header=None)
SOM_activity = pandas.read_csv(path + "TREKING/som(6,6).mean.txt", sep='\t', header=None)

edges = pandas.read_csv(path + "TREKING/edges.txt", sep='\t')
edges = pandas.DataFrame(edges, columns=['Neuron', 'Kinase Gene ID', 'Substrate Gene ID'])

G_KS = nx.Graph()

for i in range(KSIN.shape[0]):
    if KSIN['Kinase Gene ID'].iloc[i] != KSIN['Substrate Gene ID'].iloc[i]:
        G_KS.add_edge(KSIN['Kinase Gene ID'].iloc[i], KSIN['Substrate Gene ID'].iloc[i])
        
pos = nx.kamada_kawai_layout(G_KS)
node_size = 6.0
width = 0.6

matplotlib.style.use('default')
plt.rcParams["font.family"] = "Arial"
            
for time in range(1, activity_literature.shape[1]):
    
    fig, (ax0, ax1, ax2) = plt.subplots(1, 3, figsize=(18, 6))
    
    # Literature
    
    nodes = pandas.DataFrame(columns=['Gene ID', 'Functionality'])
    for kinase_index in range(1, activity_literature.shape[0]):
        if activity_literature.iloc[kinase_index, time] == -1:
            nodes = nodes.append(pandas.DataFrame({'Gene ID': [activity_literature.iloc[kinase_index, 0]],
                                                  'Functionality': [-1]}), ignore_index=True)
        elif activity_literature.iloc[kinase_index, time] == 1:
            nodes = nodes.append(pandas.DataFrame({'Gene ID': [activity_literature.iloc[kinase_index, 0]],
                                                  'Functionality': [1]}), ignore_index=True)
        elif activity_literature.iloc[kinase_index, time] == 2:
            nodes = nodes.append(pandas.DataFrame({'Gene ID': [activity_literature.iloc[kinase_index, 0]],
                                                  'Functionality': [2]}), ignore_index=True)

    node_color = ['blue' if node in nodes['Gene ID'].values and nodes['Functionality'].iloc[nodes[nodes['Gene ID'] == node].index[0]] == -1
                  else 'red' if node in nodes['Gene ID'].values and nodes['Functionality'].iloc[nodes[nodes['Gene ID'] == node].index[0]] == 1
                  else 'purple' if node in nodes['Gene ID'].values and nodes['Functionality'].iloc[nodes[nodes['Gene ID'] == node].index[0]] == 2
                  else 'lightgray' for node in G_KS]

    nx.draw(G_KS, ax=ax0, pos=pos, node_color=node_color, edge_color='lightgray', node_size=node_size, width=width, with_labels=False)

    # Temporal KiR

    nodes = pandas.DataFrame(columns=['Gene ID', 'Functionality'])
    for kinase_index in range(1, activity_KiR.shape[0]):
        if activity_KiR.iloc[kinase_index, time] == -1:
            nodes = nodes.append(pandas.DataFrame({'Gene ID': [activity_KiR.iloc[kinase_index, 0]],
                                                  'Functionality': [-1]}), ignore_index=True)
        elif activity_KiR.iloc[kinase_index, time] == 1:
            nodes = nodes.append(pandas.DataFrame({'Gene ID': [activity_KiR.iloc[kinase_index, 0]],
                                                  'Functionality': [1]}), ignore_index=True)

    node_color = ['blue' if node in nodes['Gene ID'].values and nodes['Functionality'].iloc[nodes[nodes['Gene ID'] == node].index[0]] == -1
                  else 'red' if node in nodes['Gene ID'].values and nodes['Functionality'].iloc[nodes[nodes['Gene ID'] == node].index[0]] == 1
                  else 'lightgray' for node in G_KS]

    nx.draw(G_KS, ax=ax1, pos=pos, node_color=node_color, edge_color='lightgray', node_size=node_size, width=width, with_labels=False)

    # TREKING

    edge_activity = pandas.read_csv(path + "TREKING/activity/edge_activity_composite_" + str(activity_KiR.iloc[0, time]) + "min.txt", sep='\t')
    edge_activity = pandas.DataFrame(edge_activity, columns=['Kinase Gene ID', 'Substrate Gene ID', 'Edge', 'Activity'])
    
    nodes = pandas.DataFrame(columns=['Gene ID', 'Functionality'])
    for kinase_index in range(1, activity_KiR.shape[0]):
        if activity_KiR.iloc[kinase_index, time] == -1:
            nodes = nodes.append(pandas.DataFrame({'Gene ID': [activity_KiR.iloc[kinase_index, 0]],
                                                  'Functionality': [-1]}), ignore_index=True)
        elif activity_KiR.iloc[kinase_index, time] == 1:
            nodes = nodes.append(pandas.DataFrame({'Gene ID': [activity_KiR.iloc[kinase_index, 0]],
                                                  'Functionality': [1]}), ignore_index=True)
    
    G_KS = nx.Graph()

    for i in range(KSIN.shape[0]):
        if KSIN['Kinase Gene ID'].iloc[i] != KSIN['Substrate Gene ID'].iloc[i]:
            if KSIN['Edge'].iloc[i] not in edge_activity['Edge'].values:
                G_KS.add_edge(KSIN['Kinase Gene ID'].iloc[i], KSIN['Substrate Gene ID'].iloc[i], color='lightgray', weight=0.6)
            else:
                edge_index = edge_activity[edge_activity['Edge'] == KSIN['Edge'].iloc[i]].index[0]
                if edge_activity['Activity'].iloc[edge_index] == 0:
                    G_KS.add_edge(KSIN['Kinase Gene ID'].iloc[i], KSIN['Substrate Gene ID'].iloc[i], color='lightgray', weight=0.6)
                elif edge_activity['Activity'].iloc[edge_index] == -1:
                    G_KS.add_edge(KSIN['Kinase Gene ID'].iloc[i], KSIN['Substrate Gene ID'].iloc[i], color='blue', weight=0.6)
                elif edge_activity['Activity'].iloc[edge_index] == 1:
                    G_KS.add_edge(KSIN['Kinase Gene ID'].iloc[i], KSIN['Substrate Gene ID'].iloc[i], color='red', weight=0.6)
                else:
                    G_KS.add_edge(KSIN['Kinase Gene ID'].iloc[i], KSIN['Substrate Gene ID'].iloc[i], color='purple', weight=0.6)

    node_colors = ['blue' if node in nodes['Gene ID'].values and nodes['Functionality'].iloc[nodes[nodes['Gene ID'] == node].index[0]] == -1
                  else 'red' if node in nodes['Gene ID'].values and nodes['Functionality'].iloc[nodes[nodes['Gene ID'] == node].index[0]] == 1
                  else 'lightgray' for node in G_KS]
    
    edge_colors = nx.get_edge_attributes(G_KS,'color').values()
    edge_weights = nx.get_edge_attributes(G_KS,'weight').values()

    nx.draw(G_KS, ax=ax2, pos=nx.kamada_kawai_layout(G_KS), node_color=node_colors, edge_color=edge_colors, node_size=node_size, width=list(edge_weights), with_labels=False)
    
    plt.text(x=-5.95, y=1.1, s='literature', fontsize=32.0)
    plt.text(x=-2.95, y=1.1, s='KiR', fontsize=32.0)
    plt.text(x=-0.6, y=1.1, s='TREKING', fontsize=32.0)
    plt.text(x=-6.8, y=0.8, s=str(int(activity_literature.iloc[0, time]))+" min", fontsize=24.0)
    plt.text(x=-4.1, y=0.8, s=str(int(activity_literature.iloc[0, time]))+" min", fontsize=24.0)
    plt.text(x=-1.4, y=0.8, s=str(int(activity_literature.iloc[0, time]))+" min", fontsize=24.0)
    fig.savefig(path + "combined/frames/frame-" + f"{time:03}" + ".png", format='png', dpi=300, bbox_inches='tight')

print("Done.")


# %% [code]
# All time points combined

G_KS = nx.Graph()

for i in range(KSIN.shape[0]):
    if KSIN['Kinase Gene ID'].iloc[i] != KSIN['Substrate Gene ID'].iloc[i]:
        G_KS.add_edge(KSIN['Kinase Gene ID'].iloc[i], KSIN['Substrate Gene ID'].iloc[i])
        
pos = nx.kamada_kawai_layout(G_KS)
node_size = 6.0
width = 0.6

matplotlib.style.use('default')
plt.rcParams["font.family"] = "Arial"

fig, (ax0, ax1, ax2) = plt.subplots(1, 3, figsize=(18, 6))
    
# Literature

nodes = pandas.DataFrame(activity_literature, columns=['Gene ID', 'Functionality'])

node_color = ['blue' if node in nodes['Gene ID'].values and nodes['Functionality'].iloc[nodes[nodes['Gene ID'] == node].index[0]] == -1
            else 'red' if node in nodes['Gene ID'].values and nodes['Functionality'].iloc[nodes[nodes['Gene ID'] == node].index[0]] == 1
            else 'purple' if node in nodes['Gene ID'].values and nodes['Functionality'].iloc[nodes[nodes['Gene ID'] == node].index[0]] == 2
            else 'lightgray' for node in G_KS]

nx.draw(G_KS, ax=ax0, pos=pos, node_color=node_color, edge_color='lightgray', node_size=node_size, width=width, with_labels=False)

# Temporal KiR

nodes = pandas.DataFrame(activity_KiR, columns=['Gene ID', 'Functionality'])

node_color = ['blue' if node in nodes['Gene ID'].values and nodes['Functionality'].iloc[nodes[nodes['Gene ID'] == node].index[0]] == -1
            else 'red' if node in nodes['Gene ID'].values and nodes['Functionality'].iloc[nodes[nodes['Gene ID'] == node].index[0]] == 1
            else 'purple' if node in nodes['Gene ID'].values and nodes['Functionality'].iloc[nodes[nodes['Gene ID'] == node].index[0]] == 2
            else 'lightgray' for node in G_KS]

nx.draw(G_KS, ax=ax1, pos=pos, node_color=node_color, edge_color='lightgray', node_size=node_size, width=width, with_labels=False)
    
# TREKING
    
nodes = pandas.DataFrame(activity_KiR, columns=['Gene ID', 'Functionality'])
    
G_KS = nx.Graph()

for i in range(KSIN.shape[0]):
    if KSIN['Kinase Gene ID'].iloc[i] != KSIN['Substrate Gene ID'].iloc[i]:
        if KSIN['Edge'].iloc[i] not in edges['Edge'].values:
            G_KS.add_edge(KSIN['Kinase Gene ID'].iloc[i], KSIN['Substrate Gene ID'].iloc[i], color='lightgray', weight=0.6)
        else:
            edge_index = edges[edges['Edge'] == KSIN['Edge'].iloc[i]].index[0]
            if edges['Functionality'].iloc[edge_index] == 0:
                G_KS.add_edge(KSIN['Kinase Gene ID'].iloc[i], KSIN['Substrate Gene ID'].iloc[i], color='lightgray', weight=0.6)
            elif edges['Functionality'].iloc[edge_index] == -1:
                G_KS.add_edge(KSIN['Kinase Gene ID'].iloc[i], KSIN['Substrate Gene ID'].iloc[i], color='blue', weight=0.6)
            elif edges['Functionality'].iloc[edge_index] == 1:
                G_KS.add_edge(KSIN['Kinase Gene ID'].iloc[i], KSIN['Substrate Gene ID'].iloc[i], color='red', weight=0.6)
            else:
                G_KS.add_edge(KSIN['Kinase Gene ID'].iloc[i], KSIN['Substrate Gene ID'].iloc[i], color='purple', weight=0.6)

node_colors = ['blue' if node in nodes['Gene ID'].values and nodes['Functionality'].iloc[nodes[nodes['Gene ID'] == node].index[0]] == -1
            else 'red' if node in nodes['Gene ID'].values and nodes['Functionality'].iloc[nodes[nodes['Gene ID'] == node].index[0]] == 1
            else 'purple' if node in nodes['Gene ID'].values and nodes['Functionality'].iloc[nodes[nodes['Gene ID'] == node].index[0]] == 2
            else 'lightgray' for node in G_KS]
    
edge_colors = nx.get_edge_attributes(G_KS,'color').values()
edge_weights = nx.get_edge_attributes(G_KS,'weight').values()

nx.draw(G_KS, ax=ax2, pos=nx.kamada_kawai_layout(G_KS), node_color=node_colors, edge_color=edge_colors, node_size=node_size, width=list(edge_weights), with_labels=False)
    
plt.text(x=-5.95, y=1.1, s='literature', fontsize=32.0)
plt.text(x=-2.95, y=1.1, s='KiR', fontsize=32.0)
plt.text(x=-0.6, y=1.1, s='TREKING', fontsize=32.0)
plt.text(x=-6.8, y=0.8, s='combined', fontsize=24.0)
plt.text(x=-4.1, y=0.8, s='combined', fontsize=24.0)
plt.text(x=-1.4, y=0.8, s='combined', fontsize=24.0)
fig.savefig(path + "TREKING_Thr/combined/frames/frame-166.png", format='png', dpi=300, bbox_inches='tight')

print("Done.")


# %% [markdown]
# Random models

# %% [code]
path = "..."

all_kinases_KiR = pandas.read_csv(path + "kinases.txt", sep='\t')
all_kinases_KiR = pandas.DataFrame(all_kinases_KiR, columns=['Gene ID'])

all_kinases = pandas.read_csv(path + "all_kinases.txt", sep='\t')
all_kinases = pandas.DataFrame(all_kinases, columns=['Gene ID'])

KSIN = pandas.read_csv(path + "PhosphoSitePlus_KSdata.txt", sep='\t')
KSIN = pandas.DataFrame(KSIN, columns=['Kinase Gene ID', 'Substrate Gene ID'])

KSIN_DOF = pandas.read_csv(path + "PhosphoSitePlus_KSdata_DOF.txt", sep='\t')
KSIN_DOF = pandas.DataFrame(KSIN_DOF, columns=['Gene ID', 'In degree', 'Out degree'])

data_Thr = pandas.read_csv(path + "Thr_TemporalKiR_binary.txt", sep='\t')
data_TNF = pandas.read_csv(path + "TNF_TemporalKiR_binary.txt", sep='\t')
time = data_Thr.iloc[:, 1:].columns.to_numpy(dtype=float)

validated_kinases = ['MAPK9', 'RAF1', 'BRAF', 'PRKCD', 'RPS6KB1', 'PTK2', 'MAPK3', 'MAPK1', 'CHEK1',
                    'MAPK2K1', 'MAPK2K2', 'SRC', 'MAPK11', 'MAPK12', 'MAPK13', 'MAPK14', 'PDPK1', 'AKT1']

N_iterations = 100

for iteration in range(N_iterations):
    print(iteration)

    os.makedirs(path + "Thr_RandomModels/" + str(iteration), exist_ok=False)
    os.makedirs(path + "TNF_RandomModels/" + str(iteration), exist_ok=False)

    # Randomly select kinases at each time point

    # Thr
    Thr_kinases = all_kinases_KiR
    for i in range(1, data_Thr.shape[1]):
        count_positives = (data_Thr.iloc[:, i] == 1).sum()
        count_negatives = (data_Thr.iloc[:, i] == -1).sum()
        count_zeros = all_kinases_KiR.shape[0] - (count_positives + count_negatives)

        select_kinases = np.random.choice([-1, 0, 1], all_kinases_KiR.shape[0], replace=True,
                                          p=[count_negatives/all_kinases_KiR.shape[0], count_zeros/all_kinases_KiR.shape[0], count_positives/all_kinases_KiR.shape[0]])

        Thr_kinases[data_Thr.columns[i]] = select_kinases.tolist()

    Thr_kinases = Thr_kinases.loc[~(Thr_kinases == 0).all(axis=1)]

    # TNF+Thr
    TNF_kinases = all_kinases_KiR
    for i in range(1, data_TNF.shape[1]):
        count_positives = (data_TNF.iloc[:, i] == 1).sum()
        count_negatives = (data_TNF.iloc[:, i] == -1).sum()
        count_zeros = all_kinases_KiR.shape[0] - (count_positives + count_negatives)

        select_kinases = np.random.choice([-1, 0, 1], all_kinases_KiR.shape[0], replace=True,
                                          p=[count_negatives/all_kinases_KiR.shape[0], count_zeros/all_kinases_KiR.shape[0], count_positives/all_kinases_KiR.shape[0]])

        TNF_kinases[data_TNF.columns[i]] = select_kinases.tolist()

    TNF_kinases = TNF_kinases.loc[~(TNF_kinases == 0).all(axis=1)]

    # Randomly assign kinases to neurons

    N_neurons_Thr = 36
    N_neurons_TNF = 36

    # Thr
    neurons_Thr = pandas.DataFrame(columns=['Neuron', 'Gene ID'])
    for i in range(Thr_kinases.shape[0]):
        neuron = random.randrange(N_neurons_Thr)
        neurons_Thr = neurons_Thr.append(pandas.DataFrame({'Neuron': [neuron],
                                                           'Gene ID': [Thr_kinases['Gene ID'].iloc[i]]}), ignore_index=True)

    neurons_Thr = neurons_Thr.sort_values(by='Neuron', ascending=True)

    # TNF+Thr
    neurons_TNF = pandas.DataFrame(columns=['Neuron', 'Gene ID'])
    for i in range(TNF_kinases.shape[0]):
        neuron = random.randrange(N_neurons_TNF)
        neurons_TNF = neurons_TNF.append(pandas.DataFrame({'Neuron': [neuron],
                                                           'Gene ID': [TNF_kinases['Gene ID'].iloc[i]]}), ignore_index=True)

    neurons_TNF = neurons_TNF.sort_values(by='Neuron', ascending=True)

    # Path search

    G_KS = nx.directed_configuration_model(in_degree_sequence=KSIN_DOF['In degree'].values, out_degree_sequence=KSIN_DOF['Out degree'].values, create_using=None, seed=iteration)
    G_KS = nx.DiGraph(G_KS)
    
    all_kinases_shuffle = all_kinases.sample(frac=1).reset_index(drop=True)

    # Thr
    Thr_paths = pandas.DataFrame(columns=['Neuron', 'Kinase Gene ID', 'Substrate Gene ID', 'Path Length', 'Path'])
    for neuron in neurons_Thr['Neuron'].unique():

        index = neurons_Thr[neurons_Thr['Neuron'] == neuron].index
        nodes = neurons_Thr['Gene ID'].iloc[index]

        for i in range(nodes.shape[0]):
            index_i = all_kinases_shuffle[all_kinases_shuffle['Gene ID'] == nodes.iloc[i]].index[0]
            for j in range(nodes.shape[0]):
                index_j = all_kinases_shuffle[all_kinases_shuffle['Gene ID'] == nodes.iloc[j]].index[0]

                try:
                    path_length = nx.shortest_path_length(G_KS, source=index_i, target=index_j)
                    paths = nx.shortest_path(G_KS, source=index_i, target=index_j)
                except:
                    pass
                else:
                    if path_length != 0:
                        Thr_paths = Thr_paths.append(pandas.DataFrame({'Neuron': [neuron],
                                                                    'Kinase Gene ID': [nodes.iloc[i]],  
                                                                    'Substrate Gene ID': [nodes.iloc[j]],
                                                                    'Path Length': [path_length],
                                                                    'Path': [paths]}), ignore_index=True)

    # TNF+Thr
    TNF_paths = pandas.DataFrame(columns=['Neuron', 'Kinase Gene ID', 'Substrate Gene ID', 'Path Length', 'Path'])
    for neuron in neurons_TNF['Neuron'].unique():

        index = neurons_TNF[neurons_TNF['Neuron'] == neuron].index
        nodes = neurons_TNF['Gene ID'].iloc[index]

        for i in range(nodes.shape[0]):
            index_i = all_kinases_shuffle[all_kinases_shuffle['Gene ID'] == nodes.iloc[i]].index[0]
            for j in range(nodes.shape[0]):
                index_j = all_kinases_shuffle[all_kinases_shuffle['Gene ID'] == nodes.iloc[j]].index[0]

                try:
                    path_length = nx.shortest_path_length(G_KS, source=index_i, target=index_j)
                    paths = nx.shortest_path(G_KS, source=index_i, target=index_j)
                except:
                    pass
                else:
                    if path_length != 0:
                        TNF_paths = TNF_paths.append(pandas.DataFrame({'Neuron': [neuron],
                                                                    'Kinase Gene ID': [nodes.iloc[i]],  
                                                                    'Substrate Gene ID': [nodes.iloc[j]],
                                                                    'Path Length': [path_length],
                                                                    'Path': [paths]}), ignore_index=True)

    # Time series, export mean neurons containing individual kinases

    for kinase in validated_kinases:
        kinase_index = all_kinases_shuffle[all_kinases_shuffle['Gene ID'] == kinase].index[0]
        
        # Thr
        Thr_kinase_neurons = pandas.DataFrame(columns=['Neuron'])
        for i in range(Thr_paths.shape[0]):
            if kinase_index in Thr_paths['Path'].iloc[i]:
                Thr_kinase_neurons = Thr_kinase_neurons.append(pandas.DataFrame({'Neuron': [Thr_paths['Neuron'].iloc[i]]}), ignore_index=True)
    
        Thr_kinase_neurons = DataFrame.drop_duplicates(Thr_kinase_neurons)

        Thr_TimeSeries_mean = pandas.DataFrame()
        for neuron in Thr_kinase_neurons['Neuron'].values:

            index = neurons_Thr[neurons_Thr['Neuron'] == neuron].index
            kinases = neurons_Thr['Gene ID'].iloc[index]

            kinases_indices = []
            for i in range(kinases.shape[0]):
                kinases_index = Thr_kinases[Thr_kinases['Gene ID'] == kinases.iloc[i]].index[0]
                kinases_indices = np.append(kinases_indices, kinases_index)

            kinases_TimeSeries = Thr_kinases.iloc[kinases_indices, 1:]
            kinases_TimeSeries_mean = kinases_TimeSeries.mean(axis=0)

            Thr_TimeSeries_mean = Thr_TimeSeries_mean.append(kinases_TimeSeries_mean, ignore_index=True)
            
        DataFrame.to_csv(Thr_TimeSeries_mean, path + "Thr_RandomModels/" + str(iteration) + "/" + kinase + ".txt", index=False, sep='\t', na_rep='')

        # TNF+Thr
        TNF_kinase_neurons = pandas.DataFrame(columns=['Neuron'])
        for i in range(TNF_paths.shape[0]):
            if kinase_index in TNF_paths['Path'].iloc[i]:
                TNF_kinase_neurons = TNF_kinase_neurons.append(pandas.DataFrame({'Neuron': [TNF_paths['Neuron'].iloc[i]]}), ignore_index=True)
            
        TNF_kinase_neurons = DataFrame.drop_duplicates(TNF_kinase_neurons)

        TNF_TimeSeries_mean = pandas.DataFrame()
        for neuron in TNF_kinase_neurons['Neuron'].values:

            index = neurons_TNF[neurons_TNF['Neuron'] == neuron].index
            kinases = neurons_TNF['Gene ID'].iloc[index]

            kinases_indices = []
            for i in range(kinases.shape[0]):
                kinases_index = TNF_kinases[TNF_kinases['Gene ID'] == kinases.iloc[i]].index[0]
                kinases_indices = np.append(kinases_indices, kinases_index)

            kinases_TimeSeries = TNF_kinases.iloc[kinases_indices, 1:]
            kinases_TimeSeries_mean = kinases_TimeSeries.mean(axis=0)

            TNF_TimeSeries_mean = TNF_TimeSeries_mean.append(kinases_TimeSeries_mean, ignore_index=True)
            
        DataFrame.to_csv(TNF_TimeSeries_mean, path + "TNF_RandomModels/" + str(iteration) + "/" + kinase + ".txt", index=False, sep='\t', na_rep='')

print("Done.")
```
